# Dynamic 3D chromosomal landscapes in acute leukemia

**DOI:** 10.1101/724427

**Authors:** Andreas Kloetgen, Palaniraja Thandapani, Panagiotis Ntziachristos, Yohana Ghebrechristos, Sofia Nomikou, Charalampos Lazaris, Xufeng Chen, Hai Hu, Sofia Bakogianni, Jingjing Wang, Yi Fu, Francesco Boccalatte, Hua Zhong, Elisabeth Paietta, Thomas Trimarchi, Yixing Zhu, Pieter van Vlierberghe, Giorgio G Inghirami, Timothee Lionnet, Iannis Aifantis, Aristotelis Tsirigos

## Abstract

Three-dimensional (3D) chromatin architectural changes can alter the integrity of topologically associated domains (TADs) and rewire specific enhancer-promoter interactions impacting gene expression. Recently, such alterations have been implicated in human disease, highlighting the need for a deeper understanding of their role. Here, we investigate the reorganization of chromatin architecture in T cell acute lymphoblastic leukemia (T-ALL) using primary human leukemia specimens and its dynamic responses to pharmacological agents. Systematic integration of matched *in situ* Hi-C, RNA-Seq and CTCF ChIP-Seq datasets revealed widespread changes in intra-TAD chromatin interactions and TAD boundary insulation in T-ALL. Our studies identify and focus on a TAD “fusion” event being associated with loss of CTCF-mediated insulation, enabling direct interactions between the MYC promoter and a distal super-enhancer. Moreover, our data show that small molecule inhibitors targeting either oncogenic signal transduction or epigenetic regulation reduce specific 3D interactions associated with transformation. Overall, our study highlights the impact, complexity and dynamic nature of 3D chromatin architecture in human acute leukemia.

**One Sentence Summary:** 3D chromatin alterations in T cell leukemia are accompanied by changes in insulation and oncogene expression and can be partially restored by targeted drug treatments.

## INTRODUCTION

The human genome is replete with regulatory elements such as promoters, enhancers and insulators. Recent findings have highlighted the impact of the spatial genome organization in governing the physical proximity of these elements for the precise control of gene expression ^1–3^. Genome organization is a multistep process that involves compacting chromatin into nucleosomes, chromatin fibers, compartments and into chromosome territories ^3, 4^. Multiple lines of evidence suggest that at the sub-megabase level, the genome is organized in distinct, non-overlapping regions of highly self-interacting chromatin, called topologically associated domains (TADs) ^5–7^. It is now clear that an important function of TADs is to restrict the interactions of regulatory elements to genes within the TADs, while insulating interactions of regulatory elements from neighboring domains ^3, 4^. Further evidence from our lab suggests that super-enhancers, clusters of multiple enhancers that often regulate genes that determine cellular identity or drive tumorigenesis ^8, 9^, are frequently insulated by and co-duplicated with strong TAD boundaries in cancer ^10^. TAD boundaries are enriched in binding of structural proteins (e.g. CTCF, cohesin) ^11^. Cohesin-mediated, convergently oriented CTCF-CTCF structural loops are essential for the organization of the genome into TADs ^12–14^. Recent studies have shown that abrogation of CTCF binding or inversion of its orientation in boundary regions can change TAD structure, reconfigure enhancer-promoter interactions by re-establishing loops ^15^ and lead to aberrant gene activation and developmental defects ^1, 16^.

In light of these reports, our understanding of how changes in chromatin organization contribute to cancer pathogenesis remains largely unexplored barring a few examples ^2, 17, 18^. In this study, using T cell acute lymphoblastic leukemia (T-ALL) as a model disease ^19, 20^, we investigated potential reorganization of the global chromatin architecture between primary T cell leukemia samples, leukemia cell lines and healthy T cell controls. Our analysis identified recurrent structural TAD boundary changes and significant alterations in intra-TAD chromatin interactions (TAD activity) that mirrored changes in gene expression. Both these types of alterations frequently affected effectors of oncogenic NOTCH1 signaling. As a principal example of a TAD boundary change, we identified a recurrent TAD boundary loss in T-ALL within the locus of a key driver of T cell leukemogenesis, MYC, which facilitates long-range interactions of the MYC promoter with a previously characterized NOTCH-bound super-enhancer element. Furthermore, in highlighting a direct role for NOTCH1 in organizing local chromatin architecture, inhibition of NOTCH1 signaling using gamma secretase inhibitors (γSI; a specific inhibitor of transmembrane proteolytic cleavage required for NOTCH1 receptor activation) significantly reduced chromatin looping in a number of enhancer-promoter pairs that are sensitive to γSI treatment (called “dynamic NOTCH1” sites ^21^). The loss of chromatin interactions between these enhancer-promoter loops was also associated with a significant reduction of the H3K27ac mark at the respective enhancer locus. However, a subset of enhancer-promoter loops including the MYC super-enhancer loop retained their interactions with target promoters following γSI treatment, despite being bound by dynamic NOTCH1. In exploring putative co-factors that may also be responsible for maintaining long range interactions, we identified CDK7 binding to be enriched in γSI-insensitive chromatin contacts. Pharmacological inhibition of CDK7 using the covalent inhibitor THZ1 significantly reduced MYC super-enhancer promoter contacts, underlining the complexity of factors regulating 3D architecture. Taken together, our findings provide a deeper insight into how the 3D chromatin architecture can affect the regulatory landscape of oncogenes in human leukemia and suggest that some of those changes can be reversed by targeted drug treatments.

## RESULTS

### Widespread changes in 3D chromatin landscape in human T-ALL

T-ALL accounts for approximately 25% of acute lymphoblastic leukemia cases ^22^ and is characterized by activating mutations in the transmembrane protein NOTCH1 in approximately 50% of patients ^23, 24^. *NOTCH1* mutations frequently co-occur with loss of function mutations in cell cycle regulators and epigenetic factors such as *CDKN2A* and *EZH2*, respectively ^20^. Based on gene expression signatures and flow cytometry-based immunophenotyping, T-ALL is classified into two main subtypes including the “canonical” T-ALL characterized by frequent NOTCH1 mutations with a T cell phenotype and the early T-lineage progenitor (ETP) leukemia subtype, frequently expressing stem cell and myeloid surface markers. Though the genetic drivers of T-ALL are well-characterized, it has not been investigated whether T cell transformation is associated with widespread changes in chromatin architecture. Herein, to broadly assess the global changes in chromatin architecture in T-ALL, we performed *in situ* Hi-C in nine primary peripheral blood T-ALL samples, two T-ALL cell lines (CUTLL1 ^25^ and Jurkat 26) and peripheral blood T cells from three healthy donors and integrated these datasets with CTCF binding, RNA expression changes and enhancer activity (Figure 1A). The Hi-C data were uniformly processed across all the samples using our HiC-bench platform ^27^. Quality assessment showed alignment rates that yielded a high percentage of usable long-range read pairs in all cases (Figure S1A, **Table S1)**. As an initial comparison of our Hi-C data across all the samples, we performed a Principal Component Analysis (PCA) of genome-wide “hic-ratio” insulation scores, representing the insulation capacity of every genome-wide bin, derived using the HiC-bench platform. Unsupervised clustering of the hic-ratio scores using the R package “Mclust” indicated three distinct clusters of samples, clearly separated by the first two components (Figure 1B). Cluster 1 samples were identified as mature peripheral T cells and separated from the T-ALL samples (cluster 2 and 3) by the first principal component. To independently discern the identity of clusters 2 and 3, we interrogated the expression pattern of these samples using gene signatures for canonical T-ALL and ETP-ALL derived from recent publications ^24, 28, 29^. Since we had no matched RNA-Seq for healthy T cells, we have used a publicly available set of RNA-Seq datasets on healthy T cell donors ^30^ (see also **Table S2** for external datasets). Amongst the T-ALL samples, four T-ALL samples that grouped in cluster 3 were identified to share a characteristic gene signature of the ETP-ALL subtype (Figure 1C). The expression signature of cluster 2 samples overlapped with that of canonical T-ALL, with a single exception displaying intermediate expression of both signatures. This T-ALL sample lacked canonical NOTCH1 mutation but harbored activating mutation in Interleukin 7 receptor alpha chain (*IL7R*) ^31, 32^ and deletion of *PTEN* ^33^ **(Table S3)**. T cells had no discernable expression pattern in either signature as expected. Thus, the assignment of canonical T-ALL and ETP-ALL using gene expression information explains the Hi-C insulation score variation between clusters 2 and 3 (Figure 1D). To further confirm the heterogeneity of the T-ALL samples from the Hi-C data, we calculated matrix-wide stratum-adjusted correlation coefficients using HiCRep ^34^ between the Hi-C contact matrices of all possible pairs of samples. T cell Hi-C samples from the three individual healthy donors highly correlated with each other (Figure S1B). Similarly, all the canonical T-ALL samples showed higher correlations to each other, including cross-comparisons of primary samples versus cell lines, and lower correlations to the normal samples. The correlation among ETP-ALL samples was on average higher when compared with T-ALL samples (Figure S1B), further supporting genome-wide variations in 3D architecture between the T cells and T-ALL samples and also between the two T-ALL subtypes.

**Figure 1:**
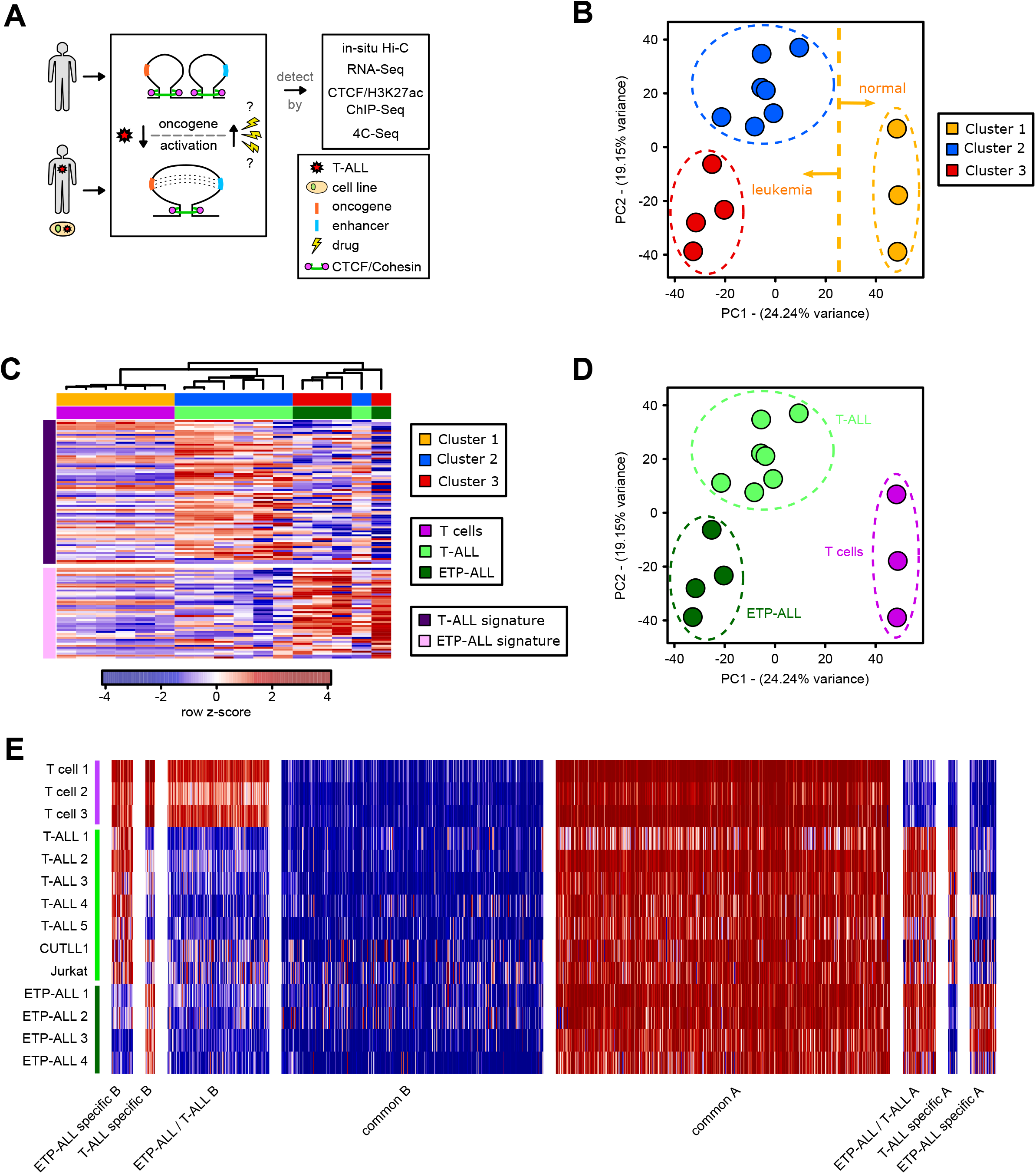
*In Situ* Hi-C analysis identifies genome-wide 3D chromatin differences between normal T cells and T-ALL subtypes. **A)** Schematic showing the overall study design. **B)** Principal Component Analysis (PCA) of the genome-wide “hic-ratio” insulation scores (defined and implemented in HiC-bench) for each Hi-C dataset identified three distinct clusters. Clustering was performed using R package Mclust, with EII and VII models showing an optimal separation using three clusters. **C)** Heatmap representation of RNA-Seq results for three clusters separated by T-ALL and ETP-ALL signature (rows). Gene signature was derived from RNA-Seq results from ^24, 28, 29^. Heatmap shows row z-score of FPKM normalized read-counts using edgeR function rpkm. **D)** Principal Component Analysis (PCA) of the genome-wide “hic-ratio” insulation scores as in B), colored by cell type assignment with the help of RNA-Seq. **E)** Compartment analysis using the c-score tool. Bins were assigned to A compartment with an average c-score > 0.1, B compartment was assigned with an average c-score < −0.1, using representative H3K27ac ChIP-Seq data for directionality (higher enrichment with H3K27ac ChIP-Seq peaks determines active compartment). Different categories of disease-specific / common compartment switches were identified using unpaired two-sided t-test on c-scores between T-ALL, ETP-ALL and T cells, and significant differences were identified using p-value < 0.1.

To better characterize the individual differences in 3D architecture that underlie this separation of the leukemic versus non-leukemic samples, we examined the compartmentalization of the genome between the three clusters of Hi-C samples. To this end, we utilized the c-score tool ^35^ to determine compartment scores and integrated H3K27ac ChIP-Seq data for CUTLL1 (T-ALL), Loucy (ETP-ALL) and T cells with resulting compartment scores to assign active (A) and inactive (B) compartments. First, we performed a PCA on genome-wide compartment scores, which showed a similar separation of T cells, T-ALL and ETP-ALL (Figure S1C) as observed before with the genome-wide insulation scores. We further identified compartment shifts that are common to both T-ALL sub-types when compared with T cells (411 A to B; 134 B to A), as well as smaller sets of compartment shifts unique to each T-ALL (40 A to B; 39 B to A) and ETP-ALL (87 A to B; 108 B to A), again highlighting the inter-sample variations among T-ALL subtypes (Figures 1E, S1D). Collectively, these data show that changes in 3D chromosomal landscapes can occur in transformed leukemia cells and can help differentiate between related subtypes of human leukemia.

### Intra-TAD activity changes affect downstream effectors of T-ALL pathogenesis

We then focused on all common TADs between T cells and T-ALL that are found within the transcriptionally active A compartment in either T cells or T-ALL. We defined the “intra-TAD activity” as the average of all normalized interaction scores of all interactions within the particular TAD (see Methods for details). Differences in the intra-TAD activity score across the selected TADs were determined by comparing the fold-change of average intra-TAD activity (as described above) between T cells and T-ALL as well as a paired t-test per interaction-bin per TAD followed by multiple testing correction. This approach approximates regulatory enhancer-promoter loops and structural loops that may discern transcriptional activity of each TAD between T cells and T-ALL. The comparison of intra-TAD activity between canonical T-ALL samples and controls identified several statistically significant gains and losses in T-ALL (Figure 2A; FDR < 0.1, TAD activity log2 fold-change > 0.58 / fold-change > 1.5). As a negative control, we performed the same comparison between two independent T cell samples, which revealed no changes by applying the same thresholds (Figure 2B; FDR < 0.1, TAD activity log2 fold-change > 0.58). Furthermore, the observed TAD activity changes were highly similar across all the T-ALL samples (Figure 2C), with some expected heterogeneity between tumors. In order to rule out that these changes were merely influenced by shifts from an active to inactive compartment or *vice versa*, we have integrated compartment shifts that overlap our differentially active TADs. We found that only ∼15-18% of the identified intra-TAD activity changes can be explained by concomitant compartment shifts, with the majority falling in the A compartment in both T cells and T-ALL samples, thus underlying a different mechanism than compartment regulation (Figure S2A).

**Figure 2:**
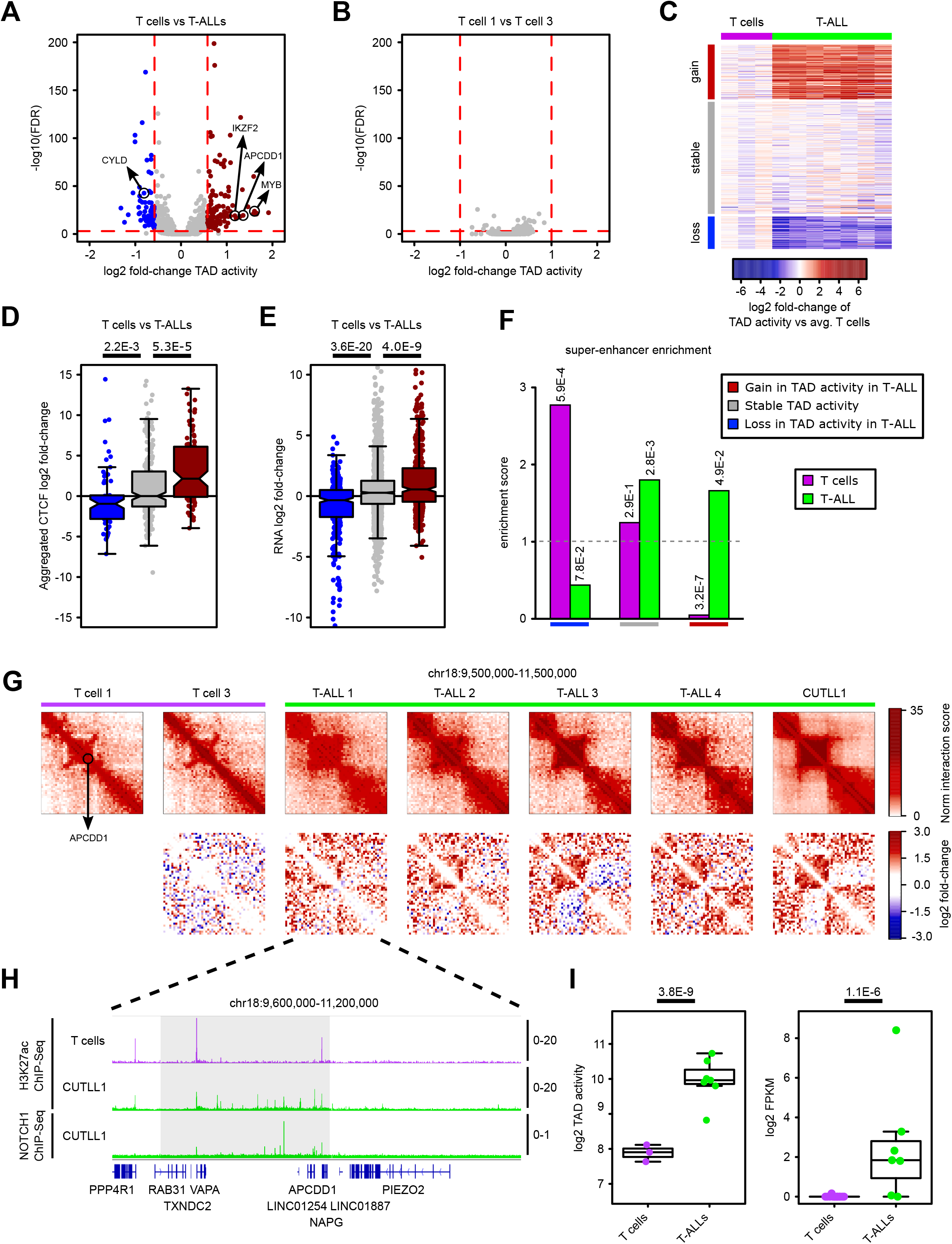
Intra-TAD activity changes affect downstream effectors of T-ALL pathogenesis. **A)** Volcano plot showing differential intra-TAD activity for pair-wise comparisons of T cells versus T-ALL. Differentially active TADs were selected using log2 fold-change of average intra-TAD activity > 0.58 or < −0.58 with FDR < 0.1 (horizontal red dotted line), and are highlighted red / blue, respectively. Statistical evaluation of each TAD was performed using paired two-sided t-test for each interaction-bin per TAD between averages of T cells and T-ALL. **B)** Volcano plot of the same analysis as in A) between two healthy T cell Hi-C samples. **C)** Heatmap showing average per-sample intra-TAD activity in all T-ALL samples and T cells normalized by the average TAD activity across all three T cell samples. Rows are showing differentially active / stable TADs as highlighted in A). **D)** Integration of CTCF binding information with TAD boundary categories from A). All CTCF bindings from surrounding TAD boundaries are aggregated, and the log2 fold-change of such CTCF signals between T-ALL and T cell is shown. Significant differences are calculated by an unpaired one-sided t-test comparing aggregated CTCF levels from TADs with decreased / increased intra-TAD activity with aggregated CTCF levels from stable TADs, assuming a positive correlation between CTCF binding and intra-TAD activity. **E)** Integration of RNA-Seq (minimum per-gene expression filter FPKM > 1) within TADs with decreased / increased intra-TAD activity. For each such gene, the respective log2 fold-change in expression between T cells and T-ALL taken from RNA-Seq is shown. Significant global differences are calculated by an unpaired one-sided t-test comparing genes from TADs with decreased / increased intra-TAD activity with genes from stable TADs, assuming a positive correlation between expression and intra-TAD activity changes. **F)** Super-enhancer integration with differentially active TADs. Only super-enhancers found mutually exclusive in either T cells or T-ALL were used. Enrichment score was calculated as observed overlap between super-enhancers and differentially active / stable TADs over expected background (cell-type specific super-enhancers in all genome-wide TADs). Statistical enrichment compared to background (cell-type specific super-enhancers in all genome-wide TADs) was calculated using two-sided Fisher’s exact test. **G)** Hi-C interaction heatmaps (first row) showing the *APCDD1* containing TAD (black rectangles). Second row shows heatmaps of per-bin log2 fold-change interactions when compared to T cell 1. **H)** H3K27ac and NOTCH1 ChIP-Seq tracks for the APCDD1 locus, shown as fold-enrichment over input. **I)** Quantifications for intra-TAD activity (left; as highlighted in G)) and expression of APCDD1 (right). Statistical evaluation for intra-TAD activity was performed using paired two-sided t-test of average per interaction-bin for APCDD1 TAD between T cells and T-ALL, followed by multiple testing correction (see methods). APCDD1 expression was determined by RNA-Seq and shown as log2 FPKM for T cells and T-ALL samples; normalization and statistical evaluation was performed using edgeR followed by multiple testing correction.

Because cancer genomes often show mutational aberrations and copy-number variants (CNVs), we investigated the impact of such aberrations on our intra-TAD activity analysis. We used HiCnv ^36^, a tool to detect potential CNVs reliably solely from Hi-C data. Following this approach, we only found between zero and nine CNVs per T-ALL sample (Figure S2B). To map genetic alterations more precisely, we performed whole-genome sequencing in two selected T-ALL samples and called CNVs and tandem-duplications genome-wide. We overlapped CNVs (separated by gain/loss vs the linear genome) and tandem duplications with increased/decreased intra-TAD activity, respectively. We found no CNV/tandem duplication within any reported differentially active TAD (Figure S2C). Furthermore, no CNV/tandem duplication occurred within the TAD boundaries adjacent to differentially active TADs, which might have influenced CTCF binding (Figure S2D). We were also interested whether single nucleotide variants (SNVs) detected from WGS are enriched in any differential TAD-activity category. Indeed, for both T-ALL samples profiled we found a minor but significant increase in SNVs per Mb in gained intra-TAD activity in T-ALL when compared to stably active TADs (Figure S2E), which could potentially lead to enhancer/transcriptional upregulation and impact the 3D interactions within TADs. Thus, we conclude that our analysis is not impacted by CNVs but to a modest degree by SNVs, with the majority of 3D chromatin alterations potentially being epigenetically regulated.

To further characterize the identified differential intra-TAD activity, we integrated genome-wide binding of the insulator protein CTCF (ChIP-Seq) from T cells and leukemic samples with our Hi-C datasets. Interestingly, the changes in intra-TAD activity strongly correlated with changes in the binding of CTCF at the boundaries of the differentially active TADs. Thus, a stronger insulation by CTCF is associated with stronger intra-TAD activity (Figure 2D), suggesting that highly active TADs are strongly insulated from adjacent TADs. Next, to investigate whether these CTCF binding-associated changes in intra-TAD interactions are also associated with changes in gene expression, we performed differential expression analysis (canonical T-ALL vs. normal) and integrated the results with the differentially active TADs. To this end, we extracted all expressed genes (FPKM > 1) falling into differentially/stably active TADs and calculated gene expression fold-changes between the leukemic sample and their normal counterparts. Following the hypothesis of a positive correlation between intra-TAD activity and gene expression, we observed that increased chromatin interactions in T-ALL significantly associated with positive fold-changes in gene expression, whereas decreased intra-TAD activity in T-ALL correlated with negative fold-changes in gene expression when compared to gene expression changes within stable TADs (Figure 2E). We then overlapped these highly correlating changes of intra-TAD activity, CTCF insulation and gene expression with cell-type specific super-enhancer activity in T-ALL and T cells. We defined super-enhancers for naïve T cells and the T-ALL cell-line CUTLL1 with the ROSE algorithm ^9^ applied on H3K27ac ChIP-Seq data and identified cell-type specific super-enhancers. We found a significant enrichment of T-ALL specific super-enhancers in the TADs that gained activity in T-ALL and, *vice versa*, a significant enrichment of T cell specific super-enhancers in TADs that lost activity (Figure 2F). Taken together, these results demonstrate the existence of a number of changes in intra-TAD activity in T-ALL cells and associated TAD activity alterations, CTCF binding, mRNA expression and super-enhancer activity.

To investigate the impact of genetic alterations on CTCF binding, we interrogated again the WGS from two primary T-ALL samples. We found that genome-wide differential CTCF binding was not particularly impacted by SNVs overlapping with CTCF binding motifs when compared to stable CTCF binding (Figure S2F). We have observed small variances when selectively comparing differential CTCF binding overlapping with SNVs within TAD boundaries of differentially active TADs. However, the majority of differential CTCF binding within altered TAD boundaries was not affected by SNVs (Figure S2G).

Our comparison of changes in TAD activity and super-enhancer firing suggests that 3D chromosomal changes should occur in loci that are important for T-ALL pathogenesis, including genes that are NOTCH1 targets and over-expressed in T-ALL samples. One such gene is the adenomatous polyposis coli downregulated 1 (APCDD1), a membrane bound glycoprotein that is overexpressed in T-ALL patient samples and is a NOTCH1 target gene that is significantly downregulated following inhibition of NOTCH1 signaling (dynamic NOTCH1 target) by γSI ^21^. Our Hi-C data showed that APCDD1 was present in a TAD that gained activity in T-ALL relative to control T cells (Figure 2G, I). The increased TAD activity was common among all the T-ALL samples, concomitant with widespread activation of enhancer elements specific to T-ALL (Figure 2H). The gain of TAD activity also correlated with increased expression of APCDD1 in T-ALL samples relative to control T cells (Figure 2I). Another example of a T-ALL-specific increase in intra-TAD activity, enhancer activity and gene expression is Ikaros family gene IKZF2 (Helios). IKZF2 overexpression in hematopoietic progenitors results in T cell developmental arrest ^37^. We were able to identify a T-ALL specific super-enhancer within the same TAD, as well as a significantly increased gene expression in T-ALL compared to normal T cells (Figure S3A, B, C). In contrast, among the TADs that lost activity in T-ALL, we identified the gene CYLD, a deubiquitinating enzyme, and a known repressor of NF-kB signaling and a putative tumor suppressor in T-ALL. Previous work from our lab and others identified that constitutive activation of NOTCH1 triggers the NF-kB pathway in T-ALL and the mechanism of NOTCH1 dependent activation of NF-kB signaling is through the repression of CYLD ^38, 39^. CYLD is a negative regulator of IKK, which promotes NF-kB signaling. We found significant loss of interactions in the TAD that harbors CYLD in all profiled T-ALL samples (Figure S3D, E). The loss of TAD activity also correlated with the decreased expression of CYLD in T-ALL samples (Figure S3F).

### Intra-TAD activity distinguishes between T-ALL subtypes

Finally, to investigate subtype specific differences in TAD activity, we evaluated the intra-TAD differences between the canonical T-ALL and ETP-ALL samples. We performed both individual comparisons of T-ALL and ETP-ALL versus untransformed T cells, and also directly compared the intra-TAD activity between T-ALL and ETP-ALL. These comparisons identified both common changes in T-ALL and ETP-ALL when compared to T cells, but also disease-specific alterations that reflect both the common lineage but also different stages of maturation arrest of the two subtypes (Figure S4A, B). Integration of gene expression changes with differentially active TADs again indicated significant correlations of intra-TAD activity changes with expression changes between ETP-ALL and T cells (Figure S4C). Similarly, we found significant correlation of expression changes with intra-TAD activity changes between ETP-ALL and canonical T-ALL (Figure S4D), highlighting the impact of 3D architecture on gene expression changes between T-ALL subtypes.

### Identification of recurrent TAD insulation changes in T-ALL

Following the identification of differences in intra-TAD activity, we further investigated TAD boundary changes between normal T cells and T-ALL samples. We performed a global TAD insulation alteration analysis on all pairs of adjacent TADs found in T cells (revealing TAD boundary losses) and, *vice versa*, on all pairs of adjacent TADs found in T-ALLs (revealing TAD boundary gains) (Figure 3A). A TAD boundary loss was defined as an increase in inter-TAD interactions accompanied by intra-TAD changes as well as loss of CTCF binding between two adjacent T cell TADs leading to a TAD “fusion” event. Conversely, a TAD boundary gain was defined as the formation of two distinct TADs (TAD “separation”) with decreased inter-TAD interactions, changes in intra-TAD interactions as well as increased CTCF binding in T-ALLs (Figure 3A) (see Methods for details). TAD fusion or separation events can change local regulatory landscapes by altering insulation of genes from nearby regulatory elements. In order to estimate a false discovery rate for our findings, we have performed the analysis first between all pairwise T cell comparisons as well as between T cells and T-ALL without integrating CTCF information (Figure 3B). Importantly, the pairwise T cell comparisons (obtained from different donors) identified only a few TAD boundary alterations between each other that could be due to the potential heterogeneity between the individual donors. However, if we consider all such insulation changes between the T cell samples as false-positives, we estimated an approximate 10.77% false discovery rate (FDR) for TAD boundary changes in T-ALL compared to T cells. Next, we integrated CTCF binding information for the T-ALL versus T cell comparisons (Figures 3C, D). To further investigate the heterogeneity of T-ALL samples for such TAD reorganization events and as an independent validation, we calculated the hic-ratio insulation score for all TAD boundary alterations found between T-ALLs and T cells. The hic-ratio score only acts as a local approximation of insulation and may miss long-range interaction changes of large TADs as interrogated specifically in our TAD boundary alteration analysis. The hic-ratio insulation score was on average significantly increased/decreased for TAD boundary gains/losses, respectively, across all T-ALL samples (Figure 3E, F).

**Figure 3:**
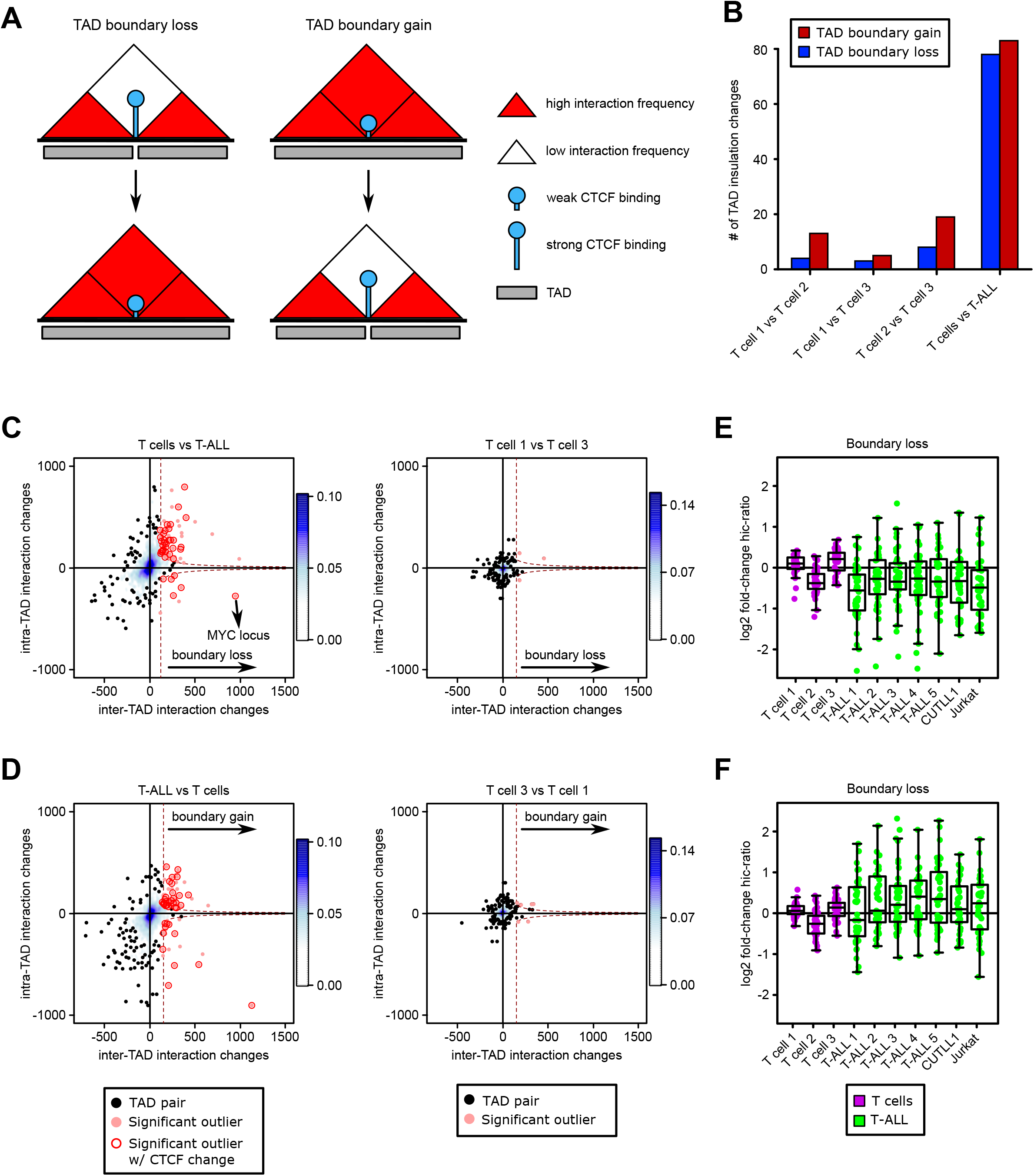
TAD boundary insulation analysis reveals changes in insulation of neighboring TADs. **A)** Schematic describing TAD boundary insulation alteration events. A TAD boundary loss (left) is associated with a strong increase in inter-TAD interactions of two adjacent T cell TADs and loss of CTCF binding. A TAD boundary gain (right) is associated with a strong decrease in inter-TAD interactions of two adjacent T-ALL TADs and concomitant gain of CTCF binding. **B)** Total numbers of TAD boundary gains / losses identified between T-ALL and T cells before integrating differential CTCF binding information. All pairwise comparisons of T cells act as negative controls. **C+D)** Representation of TAD insulation alteration events (red dots) among all pairs of adjacent TADs (black dots). Plots depict comparisons for TAD boundary losses of adjacent T cell TADs within T-ALL samples (C left), or between T cell samples 1 and 3 (C right). Plots in D) depict comparisons for TAD boundary gains of adjacent T-ALL TADs when compared to T cell samples (D left), or between T cell samples 1 and 3 (D right). Encircled adjacent TADs demarcate gain / loss of insulation accompanied by more than one gained / lost CTCF binding, respectively. Significant changes in CTCF binding were calculated using the R package DiffBind and filtered for FDR < 0.1 and log2 fold-change > 1 / < −1. **E+F)** All TAD boundary alterations (boundary loss (E), boundary gain (F)) from comparisons in C) and D) between T-ALL and T cells were used to estimate heterogeneity in insulation changes of such boundary alterations across all analyzed Hi-C samples. Hic-ratio insulation scores for each boundary and sample were compared vs. the average hic-ratio insulation score of all T cell samples. Boundary losses (*n*=78) come with a decrease in insulation scores on average, while boundary gains (*n*=83) come with increase in insulation scores across all T-ALLs on average when compared to the average hic-ratio insulation score of all T cell samples.

Lastly, using WGS data, we found only two TAD boundaries displaying increased/decreased insulation capacity in T-ALL overlapping with either genomic deletions or insertions, however, none of the indels was directly overlapping with CTCF binding motifs of differential CTCF binding sites (Figure S5A). We furthermore identified two genomic inversions (potentially leading to aberrant CTCF orientation as previously reported ^15, 40^) that overlapped reported TAD boundary insulation alterations (Figure S5B). However, none of them affects genomic loci that contain relevant T-ALL genes.

### A recurrent TAD fusion event permits *MYC* promoter/super-enhancer looping

MYC is widely up-regulated in T-ALL and is one of the main oncogenes activated downstream of NOTCH1 signaling, contributing to metabolic rewiring, cell growth and proliferation ^41, 42^. Intriguingly, we identified a recurrent TAD fusion event in the MYC locus in all T-ALL samples studied (both primary and cell lines) compared to untransformed T cells (Figure 4A). The TAD fusion event in all T-ALL samples led to a strong increase in inter-TAD interactions compared to control T cells. We then investigated whether this observed increase in inter-TAD interactions and the fusion of the two TADs into one larger TAD is a result of loss of CTCF-mediated insulation at the T cell specific TAD boundary. CTCF ChIP-Seq and ChIP-qPCR data confirmed CTCF binding at the TAD boundary in normal T cells, and an almost complete abrogation of CTCF binding at this boundary across the T-ALL samples (Figure 4B, S6A). This loss of CTCF binding was not due to mutation of the CTCF binding site in T-ALL (Figure S6B) or DNA methylation changes within the CTCF binding site countering CTCF binding in T-ALL (data not shown). Furthermore, 5-azacytidine treatment leading to global DNA de-methylation showed no restoration of CTCF binding in CUTLL1 cells (Figure S6C). Instead, ATAC-Seq data indicated a significant decrease in chromatin accessibility of the CTCF binding site in T-ALL (Figures 4B, S6D).

**Figure 4:**
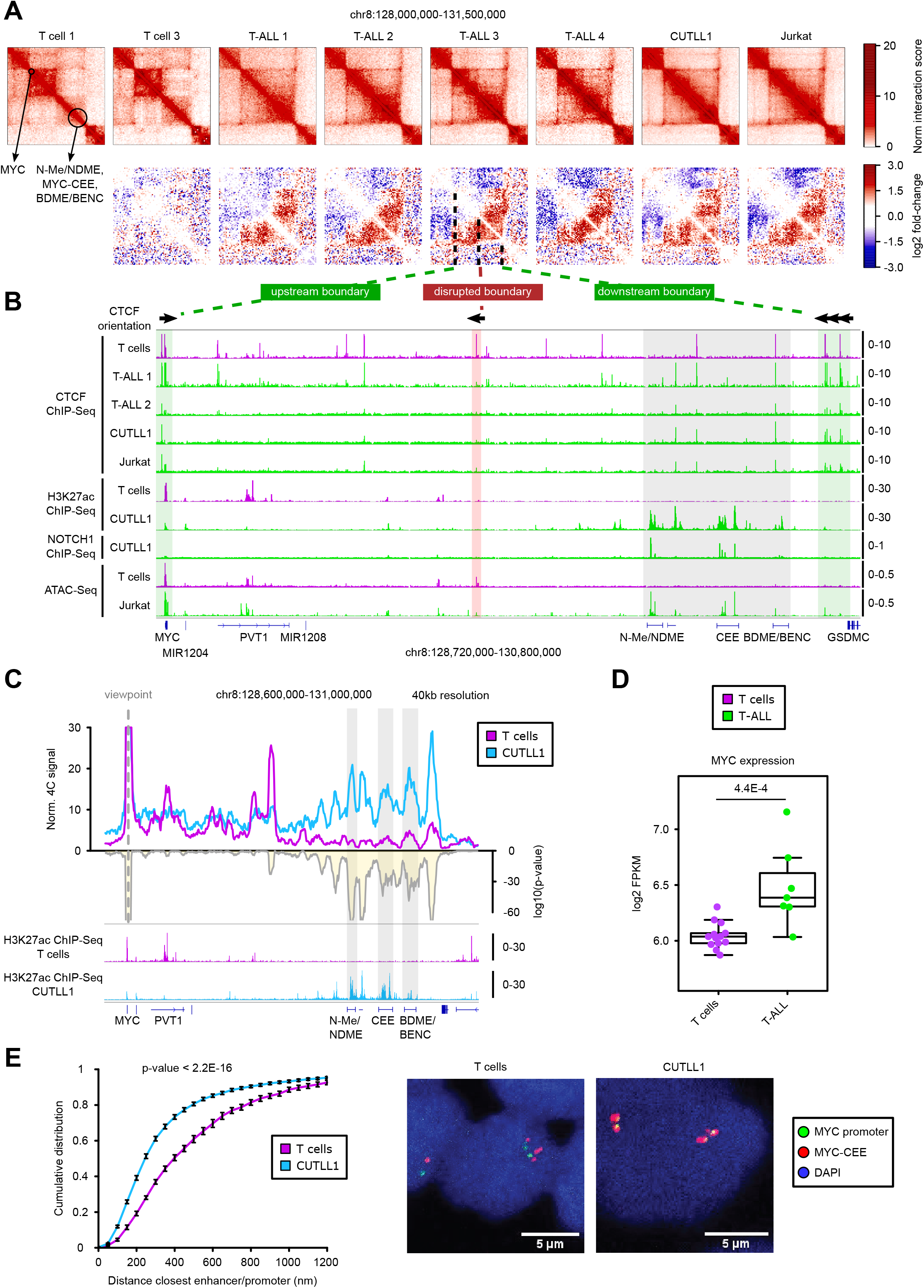
MYC overexpression in leukemia is associated with CTCF loss and TAD boundary insulation loss. **A)** Hi-C interaction heatmaps (first row) showing the MYC locus. Second row shows heatmaps of per-bin log2 fold-change interactions when compared to T cell 1. In T cells, MYC is located in the upstream TAD at its upstream boundary, while the super-enhancer cluster is located in the downstream TAD. **B)** CTCF and H3K27ac ChIP-Seq tracks for the MYC locus. CTCF orientation is shown for canonical CTCF binding motifs derived from PWMScan ^66^ (database JASPAR CORE vertebrates; filtered by p-value < 1E-5). Green boxes on the left and right highlight CTCF binding in the upstream and downstream TAD boundaries, with the downstream boundary showing multiple CTCF binding sites with the same orientation. Red box in the middle highlights loss of CTCF insulation in leukemia. Grey box towards the right highlights the super-enhancer cluster. ChIP-Seq tracks show fold-enrichment over input where applicable, counts-per-million reads otherwise. ATAC-Seq tracks show counts-per-million reads. **C)** 4C-Seq analysis using *MYC* promoter as viewpoint. Positive y-axis shows interactions with the *MYC* promoter viewpoint as normalized read counts, while negative y-axis shows significance of differential interactions between T cells and CUTLL1 as log(p-value) derived using edgeR function glmQLFTest. The three grey boxes highlight three areas of strong H3K27ac signal within the super-enhancer element (N-Me/NDME, CEE, BDME/BENC) that correlate with *MYC* promoter interactions. Tracks below show H3K27ac ChIP-Seq tracks for T cells and CUTLL1 as fold-enrichment over input. **D)** MYC expression determined by RNA-Seq and shown as log2 FPKM for T cells and T-ALL samples. Statistical evaluation was performed using two-sided edgeR analysis followed by multiple testing correction. **E)** Distance between *MYC* promoter and center enhancer element (MYC-CCE) measured by DNA-FISH analysis (left). Statistical difference between distributions of probe distances was calculated using two-sample one-sided Kolmogorov Smirnov test following the hypothesis of increased probe-distance in T cells when compared to T-ALL. Probe-pairs T cells = 993; Probe-pairs CUTLL1 = 2001. Median distance T cells = 412.84μm. Median distance CUTLL1 = 264.28μm.

In T-ALL, MYC transcription is controlled by distant 3D interactions with a long stretch of enhancers, including the previously characterized N-Me/NDME element ^43, 44^. This particular enhancer element is strongly bound by NOTCH1 (Figure 4B). As a result of the here reported TAD fusion event, the *MYC* promoter and its super-enhancer, that are separated by strong insulation in T cells, are now in close spatial proximity within the same TAD in leukemic samples (Figure 4A, B). To further validate whether this TAD boundary change and resulting TAD fusion increases interactions of the *MYC* promoter with the distal super-enhancer at high resolution, we performed 4C-Seq analysis using the *MYC* promoter as viewpoint in the leukemia samples and normal T cells. 4C-Seq confirmed the interactions between the *MYC* promoter and the super-enhancer element in primary T-ALL samples and CUTLL1, whereas in untransformed T cells, no such interaction was observed (Figure 4C, S7A). This was further highlighted by the strong insulation at the lost TAD boundary provided by the binding of CTCF in T cells. Interestingly, our analysis showed that the strongest and most significant interactions specifically overlap with H3K27ac ChIP-seq peaks throughout the entire super-enhancer element, including an uncharacterized putative center enhancer element (from here on termed MYC-CEE) and the BDME/BENC enhancer cluster recently identified to be essential for normal hematopoiesis and AML pathogenesis (Figure 4C, S7A) ^45, 46^. In agreement with our 3D chromosomal interaction data, MYC was overexpressed in our T-ALL cohort compared to T cells from healthy donors (Figure 4D). Furthermore, we independently validated the differential 3D localization of *MYC* and the super-enhancer between T cells and T-ALL using 3D Fluorescence in situ hybridization (FISH) with probes targeting the *MYC* promoter and the center super-enhancer element MYC-CEE. Inter-probe distance was significantly higher in T cells compared to T-ALL cells (the CUTLL1 cell line) consistent with the increased interactions in CUTLL1 identified by 4C-Seq (Figure 4E). To confirm the CTCF-mediated insulation of the *MYC* TAD in T cells, we disrupted the CTCF binding site in normal T cells using CRISPR (clustered regularly interspaced short palindromic repeats) mutation (Figure S8A). We achieved ∼92% of cells harboring indels of varying sizes located within the CTCF motif (Figure S8B). Mutations of the CTCF motif in T cells resulted in significantly decreased CTCF binding in the edited T cells (Figure S8C) and marginally increased MYC expression (Figure S8D). The decreased binding of CTCF protein was accompanied by significantly reduced interactions between the *MYC* promoter and the CTCF bound TAD boundary region in edited T cells compared to WT T cells (Figure S8E).

### Pharmacologic NOTCH1 inhibition leads to a decrease of 3D interactions in a group of NOTCH1-regulated loci

Our analysis revealed widespread changes in global TAD structure and intra-TAD activity affecting important genes in T-ALL. However, the question whether oncogenic drivers, such as NOTCH1, play a direct role in these changes and whether their inhibition can reverse these changes remains open. To address this, we performed *in situ* Hi-C analysis in CUTLL1 cells treated with the NOTCH pathway inhibitor targeting the gamma secretase complex (γSI) for 72h as performed previously ^21, 41^. γSI selectively and efficiently inhibits NOTCH1 signaling and has strong anti-leukemic effects ^42, 47^. Hi-C analysis following γSI treatment did not reveal any significant changes either in intra-TAD activity (Figure S9A) or reversal of TAD boundary insulations (Figure S9B). This was not surprising since NOTCH1, a *bona fide* signal-dependent transcription factor, was not expected to impact global chromatin architecture. However, it was previously shown that about 90% of NOTCH1 binding sites that are sensitive to γSI treatment (i.e. dynamic NOTCH1 sites) are localized outside promoters in putative distal enhancer elements. These dynamic NOTCH1-occupied enhancers also showed significant changes in H3K27ac signal following NOTCH1 inhibition ^21^. We investigated whether chromatin interactions between dynamic NOTCH1-occupied enhancers and neighboring promoters were altered following γSI treatment. To this end, we first profiled H3K27ac following γSI treatment and categorized all non-promoter H3K27ac peaks as either stable peaks that do not change following γSI treatment or those that have either a strong reduction or increase in H3K27ac signal following γSI treatment (Figure 5A). As previously observed, the H3K27ac peaks that had a strong reduction in signal following γSI treatment were also significantly enriched for dynamic NOTCH1 binding when compared to stable or increased H3K27ac signals^21^ (Figure 5B). An increase in H3K27ac upon γSI treatment is thus likely a downstream effect of NOTCH1 inhibition. To connect NOTCH1 pathway inhibition, changes in H3K27ac and 3D looping, we used Hi-C data following γSI treatment to quantify changes in chromatin interactions of H3K27ac-enriched chromatin loops identified by H3K27ac HiChIP in CUTLL1 ^48^. Our HiChIP data showed strong enrichment of promoter-enhancer interactions as represented by a virtual 4C analysis using the MYC promoter as virtual viewpoint, as well as reliable detection of such loops from our pipeline (Figure S9C). Dynamic NOTCH1-bound enhancers with reduced H3K27ac levels following γSI treatment showed the strongest loss of chromatin interactions with connected genes (Figure 5C). Interestingly, dynamic NOTCH1-bound enhancers with no change in H3K27ac signal on average remained in stable contact with nearby promoters. To correlate these changes in chromatin interaction with the dynamics of NOTCH1-dependent transcription, we also performed global run of sequencing (GRO-Seq)^49^ to measure nascent transcription following NOTCH1 inhibition by γSI and release after inhibitor “wash off” for increasing time intervals. Interestingly, the regulatory enhancer-promoter contacts most sensitive to γSI treatment were among genes that showed significant response in transcription to NOTCH1 inhibition and release after γSI wash off (Figure 5D).

**Figure 5:**
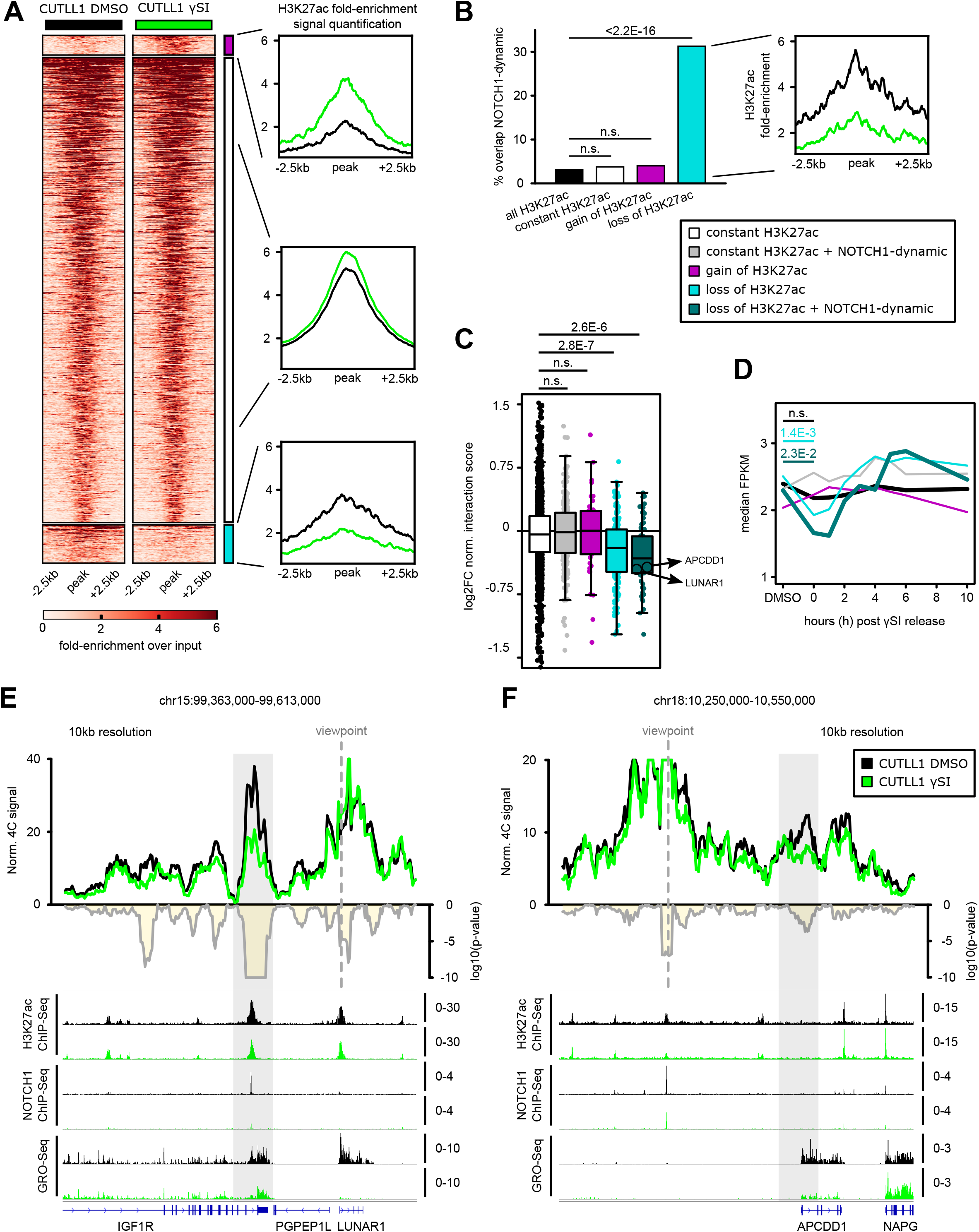
NOTCH1 inhibition affects promoter-enhancer looping specifically of NOTCH1-dependent enhancers. **A)** Differential H3K27ac occupancy analysis based on H3K27ac ChIP-Seq in CUTLL1 with and without NOTCH1-inhibitor γSI. The three identified groups consist of stable non-promoter H3K27ac signal (middle, black, *n*= 2949), increased (upper, pink, *n*=125) and reduced non-promoter H3K27ac signal (lower, light-blue, *n*=243). Heatmap shows the H3K27ac signal as fold-enrichment over input within +/− 2.5kb around the summit of the identified H3K27ac peaks and line plots depict quantification of H3K27ac signal within these regions, both created with DeepTools ^67^. Differential analysis was performed with the R package DiffBind and differential peaks were selected using FDR < 0.05, log2 fold-change > 1.0 or < −1.0. **B)** Overlap of constant, increased and reduced H3K27ac peaks with NOTCH1-dynamic sites previously defined by ^21^. Quantification of H3K27ac signal shown as fold-enrichment over input (right panel) specifically for peaks with reduced H3K27ac signal and dynamic NOTCH1 binding (*n*=76). Statistical evaluation was performed using two-sided Fisher test against all non-coding H3K27ac peaks overlapping dynamic NOTCH1 binding. **C)** Changes in chromatin interactions upon γSI between non-promoter H3K27ac peaks of interest defined in A) and B) and nearby gene promoters (defined using H3K27ac HiChIP data in CUTLL1) are shown as log2 fold-change of normalized interaction score. Each dot represents such a promoter-enhancer interaction defined by H3K27ac HiChIP. Significance of global shifts compared to enhancer-promoter loops of stable enhancers (grey, left) is calculated by unpaired one-tailored t-test, following the hypothesis of a positive correlation between enhancer activity and promoter-looping. **D)** Changes in gene expression upon γSI for all genes defined in C) are shown as log2 fold-change of FPKM calculated from GRO-Seq data. Significance of global differences when compared to genes associated with stable H3K27ac signal is calculated by unpaired one-tailored t-test, following the hypothesis of a positive correlation between promoter-enhancer looping and gene expression. **E+F)** 4C-Seq analysis using *LUNAR1* promoter (E) or *APCDD1* enhancer (F) as viewpoints. Positive y-axis shows interactions with the viewpoint as normalized read counts, while negative y-axis shows significance of differential interactions between untreated and γSI treated CUTLL1 as log(p-value) calculated using edgeR function glmQLFTest. The grey box in E) shows a previously reported intronic *IGF1R* enhancer; the grey box in F) shows the *APCDD1* promoter. Tracks below show H3K27ac and NOTCH1 ChIP-Seq and GRO-Seq (positive strand only) before and after γSI treatment as fold-enrichment over input where applicable, counts-per-million otherwise.

To further validate the changes in interactions among the NOTCH1-sensitive enhancer-promoter loops, we performed 4C-Seq analysis for two previously characterized NOTCH1 T-ALL targets, LUNAR1 and APCDD1. LUNAR1 is a long non-coding RNA that we have previously identified as a *cis* regulator of the expression of the neighboring *IGF1R* gene. This regulation is achieved by chromatin looping of *LUNAR1* promoter with an intronic enhancer in *IGF1R* ^50, 51^. 4C-Seq using *LUNAR1* promoter as viewpoint identified strong interactions between *LUNAR1* promoter and the *IGF1R* enhancer element. However, the interaction decreased significantly following NOTCH1 inhibition (Figure 5E, S9D). The decrease in interaction also correlated with reduction in H3K27ac signal at the enhancer element and decreased expression of LUNAR1 (Figure 5E, S9D). Similarly, in the APCDD1 locus, 4C-Seq using an *APCDD1* enhancer with dynamic NOTCH1 and reduced H3K27ac signal as viewpoint, we identified decreased interaction between the enhancer and the promoter of *APCDD1*. These changes also correlated with reduced H3K27ac in the enhancer and reduced expression of APCDD1 (Figure 5F, S9E). These results suggest that pharmacologic NOTCH1 inhibition can affect activity (as defined by H3K27ac levels) of dynamic NOTCH1-bound enhancers and that 3D interactions in such enhancers are significantly diminished.

However, our analysis revealed that a subset of dynamic NOTCH1-regulated loci was not associated with either H3K27ac loss or reduced long-range chromatin interactions following γSI treatment. Based on this observation, we classified enhancers with reduced NOTCH1 binding and H3K27ac levels upon γSI treatment as γSI-sensitive enhancers, and enhancers with only reduced NOTCH1 binding as γSI-insensitive enhancers (Figure S9F). Interestingly, γSI-sensitive enhancers tend to be shorter in length than γSI-insensitive enhancers (Figure S9G). For example, our initial 4C analysis had identified contacts of the *MYC* promoter with three enhancer clusters. 4C-Seq analysis detected no significant decrease in the frequency of interactions between the *MYC* promoter and all of the three enhancer clusters following γSI treatment (Figure S10A, B), although γSI treatment reduced MYC expression (to approximately half by qPCR and western blot; Figure S10B, C) and NOTCH1 binding at the MYC super-enhancer (Figure S10A, B). We also noticed only moderate changes in the H3K27ac distribution within the NOTCH1-bound enhancer elements after γSI treatment (Figure S10A, B). Also, the critical CTCF binding within the TAD boundary of MYC was not restored upon γSI treatment (Figure S10D). Thus, despite the downregulated MYC mRNA expression and the loss of NOTCH1 binding, pharmacological inhibition of NOTCH1 signaling was not able to alter 3D interactions in this locus. As an additional example, a dynamic NOTCH1-bound enhancer looping to the *IKZF2* promoter did not lose interactions following γSI treatment (Figure S10E, F), suggesting that NOTCH1 binding is critical for maintaining enhancer-promoter contacts in only a subset of such loops and additional chromatin regulators may play a role in maintaining chromatin interactions of the γSI-insensitive loops.

### CDK7 inhibition targets γSI-insensitive enhancer-promoter loops

To further understand the differential sensitivity of the dynamic NOTCH1-bound enhancers, we performed a differential binding analysis using LOLA ^52^ between the γSI-sensitive and insensitive enhancers using publicly available ChIP-Seq datasets from the LOLA database. Among the chromatin regulators and transcription factors with available ChIP-Seq datasets in T-ALL cells, we found cyclin-dependent kinase 7 (CDK7) binding to be significantly enriched in the γSI-insensitive enhancers compared to the sensitive enhancers (Figure 6A). CDK7 controls the dynamics of transcriptional initiation by the phosphorylation of RNA polymerase II CTD at Ser-5 and Ser-7 ^53^. CDK7 inhibition has been particularly linked to suppression of super-enhancer linked oncogenic transcription ^54–56^. To globally assess the role of CDK7 binding in the maintenance of γSI-insensitive enhancer-promoter loops, we performed *in situ* Hi-C analysis in CUTLL1 cells treated with the CDK7 inhibitor THZ1, which was previously demonstrated to have strong anti-leukemic activity ^54^. As before, we first profiled H3K27ac levels following THZ1 treatment by ChIP-Seq and categorized all non-promoter H3K27ac peaks as either insensitive with stable H3K27ac signal, peaks that have a strong reduction (THZ1 lost enhancer) or peaks with an increase (THZ1 gained enhancer) in H3K27ac signal following CDK7 treatment (Figure 6B). Globally, as previously observed in the γSI treatment, enhancers with a strong reduction in signal of H3K27ac peaks had a correlative strong reduction in long-range chromatin interactions to target promoters, whereas the insensitive enhancers with no change in H3K27ac signal following THZ1 treatment, neither gained nor lost chromatin interactions on a global scale (Figure 6C).

**Figure 6:**
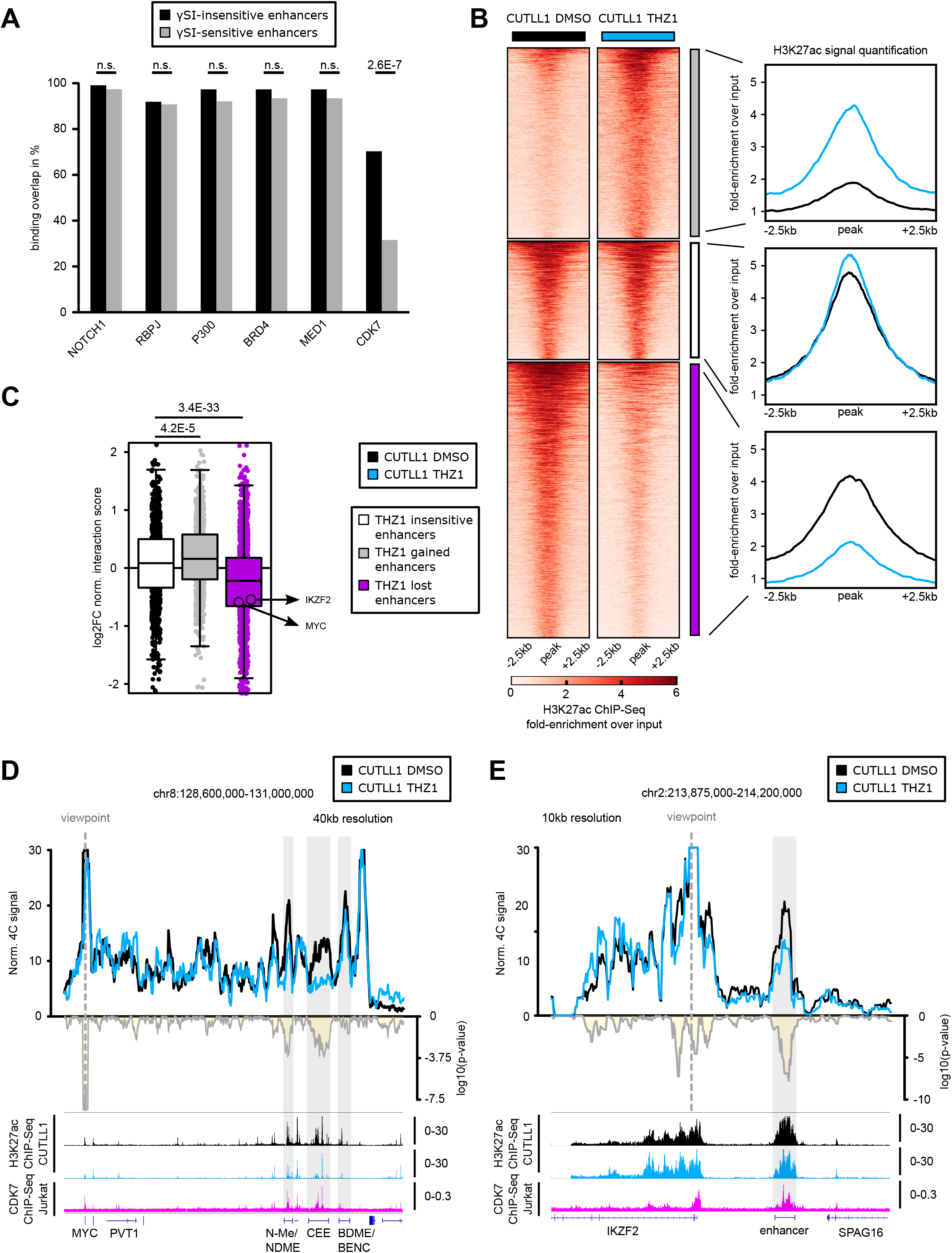
CDK7 inhibition concomitantly reduces H3K27ac levels and associated promoter-enhancer looping. **A)** LOLA analysis result for the overlap between public ChIP-Seq data in Jurkat from the LOLA database with γSI-insensitive and γSI-sensitive enhancers. Statistical differences in overlap between γSI-insensitive and sensitive enhancers with ChIP-Seq peaks were calculated using a two-sided Fisher exact test. **B)** Differential H3K27ac occupancy analysis based on H3K27ac ChIP-Seq in CUTLL1 treated with either DMSO or THZ1. The three identified groups consist of stable non-promoter H3K27ac signal (middle, white, *n*=1396), increased (upper, grey, *n*=2246) and reduced non-promoter H3K27ac signal (lower, pink, *n*=3248). Heatmap shows the H3K27ac signal as fold-enrichment over input within +/− 2.5kb around the summit of the identified H3K27ac peaks and line plots depict quantification of H3K27ac signal within these regions. Differential analysis was performed with the R package DiffBind and differential peaks were selected using FDR < 0.05, log2 fold-change > 1.0. **C)** Hi-C integration with H3K27ac peaks of interest. Changes in chromatin interactions upon THZ1 between non-promoter H3K27ac peaks of interest defined in B) and nearby gene promoters (defined using H3K27ac HiChIP data in CUTLL1) are shown as log2 fold-change of normalized interaction score. Each dot represents such a promoter-enhancer interaction. Significance of global shifts compared to enhancer-promoter interactions associated with stable enhancers (grey, left) is calculated by an unpaired one-sided t-test, following the hypothesis of a positive correlation between enhancer activity and promoter-looping. **D+E)** 4C-Seq analysis using *MYC* promoter (D) or *IKZF2* promoter (E) as viewpoints. Positive y-axis shows interactions with the viewpoint as normalized read counts, while negative y-axis shows significance of differential interactions between untreated and THZ1 treated CUTLL1 as log(p-value) calculated using edgeR function glmQLFTest. The grey boxes show the enhancer elements. Tracks below show H3K27ac before and after THZ1 treatment and CDK7 ChIP-Seq track from Jurkat cell line, and represent fold-enrichment over input where applicable and counts-per-million reads otherwise.

To further test the role of CDK7 in maintaining DNA loops, we performed high-resolution 4C-Seq following THZ1 treatment in the previously identified γSI-insensitive *MYC* and *IKZF2* loci. We observed a significant decrease in the interaction between both N-Me/NDME and MYC-CEE and the *MYC* promoter following the CDK7 treatment (Figure 6D, S11A). These changes were accompanied by a significant decrease of the H3K27ac signal and MYC expression (Figure 6D, S11A, S11B). Finally, no significant gain in the binding of CTCF to the TAD boundary was observed, suggesting that the described loss of the promoter-enhancer interaction occurs independently of CTCF binding (Figure S11C). Additionally, DNA FISH analysis confirmed a significant increase in 3D distance between the *MYC* enhancer and promoter probes following THZ1 treatment (Figure S11D). We also performed THZ1 treatment followed by 4C-Seq on the *MYC* promoter viewpoint in Jurkat cells, which display the same TAD fusion and super-enhancer activity as CUTLL1. Our results indicate a global reduction of interactions throughout the entire super-enhancer element in Jurkat (Figure S11E, F). Furthermore, similar loss of both enhancer activity and enhancer-promoter interaction was also observed in the *IKZF2* locus as shown by the H3K27ac ChIP-Seq data and 4C-Seq in CUTLL1 cells (Figure 6E, S11G). Overall, these studies demonstrate that targeting a single transcription factor (NOTCH1) is able to affect only a subset of 3D promoter-enhancer interactions associated with dynamic NOTCH1. Additional factors such as CDK7 can maintain contacts in a subset of NOTCH1-insensitive enhancers in T-ALL. Furthermore, changes in H3K27ac levels emerge as a reliable indicator of gain or loss of chromatin interactions following these drug treatments.

## DISCUSSION

Despite the intense focus on the regulatory role of TADs in human disease, it remains largely unexplored whether TAD boundary or intra-TAD activity changes are important for tumor initiation and/or maintenance. Indeed, aberrant activation of cancer drivers by enhancer hijacking remains the primary known mechanism linking 3D structural changes to oncogenic transformation ^2, 3, 57, 58^. Our studies further these findings using T-ALL as a model. They highlight the underlying complexity of factors regulating the 3D landscape in human leukemia with notable variations among different leukemia sub-types and suggest that drugs with reported anti-leukemic activity partially reverse 3D interactions in selected loci, potentially accounting for the anti-leukemogenic effects of these drugs.

Frequent loss of TAD boundary insulation has been previously observed in human cancer, including T-ALL ^58^. Consistent with these findings, we identify here a TAD boundary loss within the MYC locus that resulted in loss of insulation and enhancer hijacking. MYC is an important downstream target of NOTCH1 that activates anabolic pathways to sustain proliferation induced by constitutive NOTCH1 activation ^41, 42^. Our observations suggest that MYC upregulation in leukemic cells is associated with differences in local chromatin architecture. At this point, it is not clear what causes the loss of CTCF binding within the TAD boundary in T-ALL cells, although our preliminary studies have excluded a role for DNA methylation and somatic mutations within the CTCF motif. Interestingly, using ATAC-Seq data, we found that the CTCF site is accessible in T cells but displayed greatly reduced accessibility in T-ALL cells suggesting differential chromatin accessibility as a potential mechanism of regulating CTCF binding. In support of this hypothesis, a recent report identified chromatin accessibility around CTCF binding sites to be greatly reduced during the interphase to pro-metaphase transition ^59^. Furthermore, reduced chromatin accessibility correlated with reduced CTCF binding and loss of TAD and CTCF loop structures in pro-metaphase. In addition to the lost CTCF boundary in T-ALL, we also observed an increase in CTCF binding with the same orientation (facing into the TAD and towards MYC) downstream of the super-enhancer. Such clusters of CTCF surrounding super-enhancers have recently been described as super-anchors forming so-called stripes to ensure super-enhancer mediated regulation of nearby genes ^60^.

Besides the boundary gains and losses that affect TAD structures, we also found prevalent changes in intra-TAD activity among the TAD structures that remained constant between T-ALL and T cells. The changes in intra-TAD activity correlated with expression changes, enhancer/super-enhancer activity and insulation mediated by CTCF binding in those TAD boundaries, which appeared to be independent of compartment shifts. Supporting a prominent role for intra-TAD activity changes in modulating gene expression, recent studies tracking 3D chromatin modifications during the course of developmental processes such as embryonic stem cell differentiation and neural development did not identify major structural differences in TAD boundaries, but instead identified significant changes in interactions within TADs and sub-TADs that correlated with transcriptional levels and epigenetic states ^61, 62^. Furthermore, in line with our findings, negative correlations of intra-TAD interactions with repressive histone marks have been reported in EZH2 mutant lymphomas ^63^. Herein, our observation suggests that expression changes observed following tumor initiation are frequently associated with correlative changes in intra-TAD activity, CTCF insulation and enhancer activation.

Finally, we also addressed the direct role of oncogenic NOTCH1 in organizing the 3D chromosomal landscape associated with leukemic transformation and to what extend the identified changes can be reversed by inhibiting NOTCH-signaling. NOTCH signaling inhibition is a powerful means to inhibit leukemic growth of NOTCH1 activated T cells ^47, 64^. The effects of γSI were reported to be selective to the dynamic NOTCH1 sites, which are predominantly located within enhancers ^21, 44^. Such dynamic NOTCH1 sites are also associated with a decrease in enhancer activity following γSI treatment. These findings prompted us to further investigate the impact of NOTCH1 inhibition on the remodeling of the 3D landscape in leukemia. Our study showed that NOTCH1 inhibition using γSI had no effect on global 3D chromatin structure and intra-TAD activity, but partially targeted enhancer-promoter interactions in selected NOTCH1-regulated loci. More specifically, we identified enhancer-promoter loops of dynamic NOTCH1-bound enhancers that were also associated with a decrease in H3K27ac following γSI treatment were particularly sensitive to NOTCH1 inhibition. These results concur with a recent report that demonstrated a role for NOTCH1 in facilitating specific long-range interactions in triple-negative breast cancer and mantle cell lymphoma models ^18^.

In further understanding the differential importance for NOTCH1 binding in maintaining certain enhancer-promoter loops but not others, we found that enhancers most sensitive to NOTCH1 inhibition tend to be shorter in length. The longer stretch of the insensitive enhancers might enable other factors to bind and/or keep the chromatin in an open and accessible state for long-range chromatin interactions ^65^, thus offering a potential explanation for the variances in promoter-enhancer looping changes we observed for the NOTCH1-targets, including MYC, IKZF2, APCDD1 and LUNAR1. In agreement with this hypothesis, we found enrichment for CDK7 binding in γSI-insensitive enhancers over γSI-sensitive enhancers. CDK7 inhibition has shown significant effects in hematological malignancies and other cancer ^54–56^. Pharmacological inhibition of CDK7 by THZ1 resulted in widespread decrease in enhancer activity as quantified by H3K27ac levels. Enhancers with strong reduction of H3K27ac were also associated with correlative decrease in enhancer-promoter contacts. In turn, enhancer activity together with chromatin looping in the γSI-insensitive example loci MYC and IKZF2 have been concordantly reduced upon THZ1 treatment. This clearly highlights the complexity of super-enhancer activity and factors that dictate their interactions with gene promoters. Overall, our study underscores the need for further investigation of factors that rewire long-range interactions especially during tumorigenesis, as they could be potential targets for small molecule drug development.

## ACKNOWLEDGEMENTS

We thank all members of the Aifantis/Tsirigos laboratories for discussions throughout this project. A. Heguy and the NYU Genome Technology Center (supported in part by National Institutes of Health (NIH)/National Cancer Institute (NCI) grant P30CA016087-30) for expertise with sequencing experiments; the NYU Flow Cytometry facility for expert cell sorting; the Applied Bioinformatics Laboratory for computational assistance; Genewiz for expertise with sequencing experiments. This work has used computing resources at the High-Performance Computing Facility at the NYU Medical Center. We would also like to acknowledge Bing Ren and Anthony Schmitt for the support on the Hi-C experiments. I.A. is supported by the NIH (1P01CA229086-01A1, RO1CA216421, RO1CA202025, R01CA133379, R01CA149655, 5R01CA173636, 1R01CA194923), the Alex’s Lemonade Stand Cancer Research Foundation, the Chemotherapy Foundation, The Leukemia and Lymphoma Society and the NYSTEM program of the New York State Health Department. A.T. is supported by the American Cancer Society (RSG-15-189-01-RMC), the Leukemia and Lymphoma Society and the St. Baldrick’s Foundation. P.T. was previously supported by an AACR Incyte Corporation Leukemia Research Fellowship and is currently supported by a Young Investigator Grant from Alex’s Lemonade Stand Cancer Research Foundation. P.N. was supported by the NCI (R00CA188293) and the American Society of Hematology. P.V.V was supported by an ERC Starting Grant (639784). A.T. is a Scientific Advisor to Intelligencia.AI. All other authors declare that they have no competing financial interests.

## AUTHOR CONTRIBUTIONS

A.T. and I.A. conceived, designed and supervised the study with input from A.K., P.T. and P.N. A.K. designed and performed most of the computational analyses with help from A.T., S.N. and C.L. P.T. and P.N. designed and performed most of the experiments with help from Y.G. X.C., H.H., S.B., J.W., T.T., Y.F., F.B., Y.Z., E.P., P.V.V., G.G.I. and T.L.. P.T., P.N. and Y.G. performed Hi-C, HiChIP and 4C experiments with help from X.C., S.B., J.W. and Y.Z.. P.T. performed DNA FISH with help from Y.F. and T.L. P.T. performed ChIP-Seq with help from P.N., S.B. and J.W.. P.T. and H.H. performed RNA-Seq. T.T. performed GRO-Seq. A.T., I.A. A.K. and P.T. wrote the manuscript with input from all authors.

## DATA AVAILABILITY

All sequencing data created within this study was uploaded to NCBI GEO (https://www.ncbi.nlm.nih.gov/geo/) and is available under the accession GSE115896.

## CODE AVAILABILITY

All code for Hi-C analysis is available within the previously published Hi-C bench platform (https://github.com/NYU-BFX).

## FIGURE LEGENDS

**Supplementary Figure 1:**
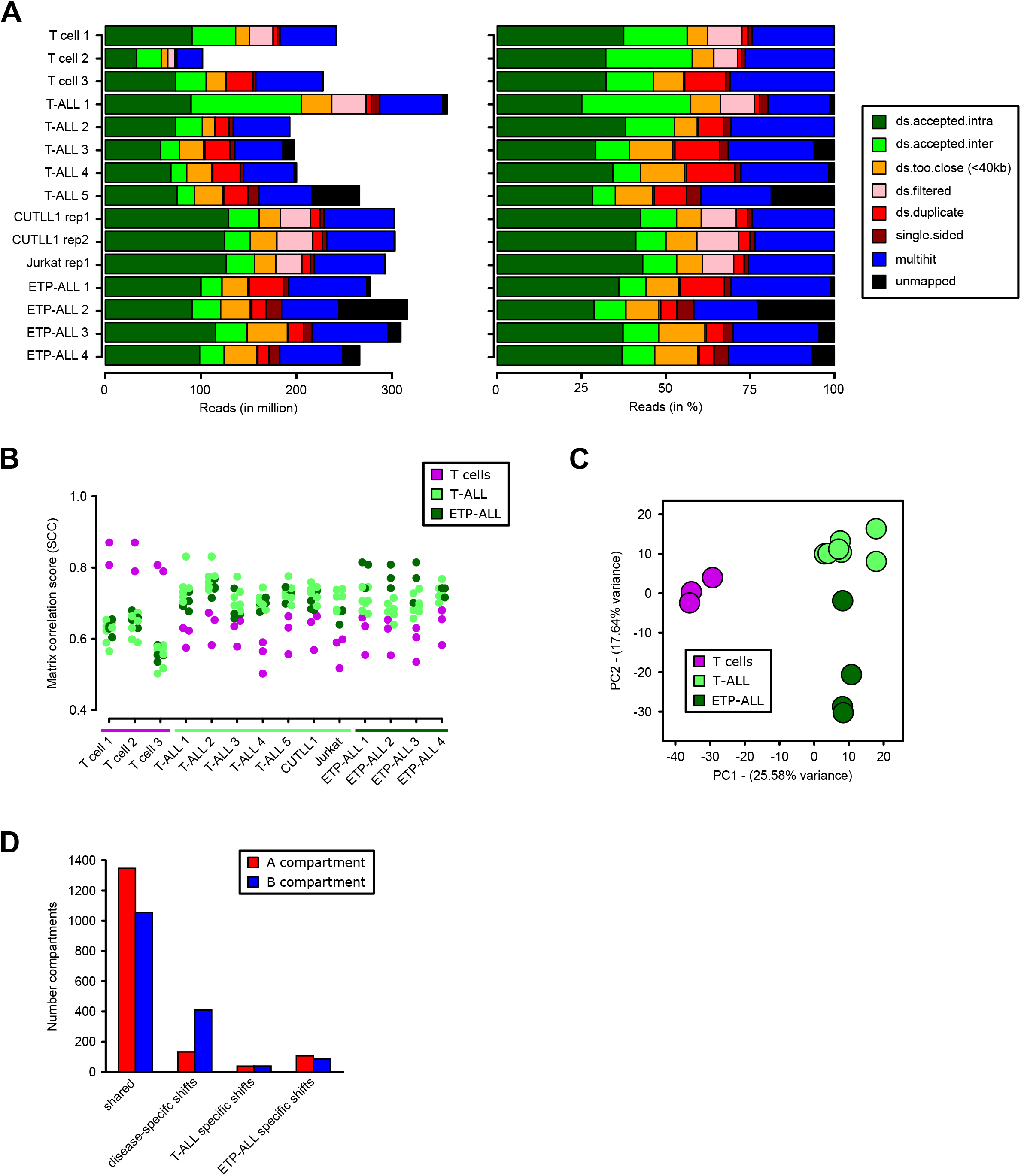
**A)** Read alignment statistics for Hi-C datasets generated within this study, as absolute reads (left) and relative reads (in %, right). “ds.accepted.intra” are all intra-chromosomal reads used for all downstream analyses. **B)** Genome-wide stratum-adjusted correlation coefficient (SCC) scores for all pair-wise comparisons of the three T cell, five canonical primary T-ALL samples, two T-ALL cell lines and four ETP-ALL Hi-C datasets. HiCRep was used to calculate chromosome-wide correlation scores, which were averaged across all chromosomes for each pair-wise comparison. The HiCRep smoothing parameter X was set to 1.0. **C)** Principal Component Analysis (PCA) of the genome-wide compartment scores for each Hi-C dataset, separating normal from leukemic samples on PC1. **D)** Compartment shifts identified between T cells, T-ALL and ETP-ALL. Assignment of A compartment was done using an average c-score > 0.1 in either all T cell, T-ALL or ETP-ALL samples as well as higher enrichment in H3K27ac signal and vice versa B compartment for average c-score < −0.1. Significance for differences between pairwise comparisons of T cells, T-ALL and ETP-ALL was determined using a two-sided t-test between c-scores, and compartment shifts were determined using p-value < 0.1.

**Supplementary Figure 2:**
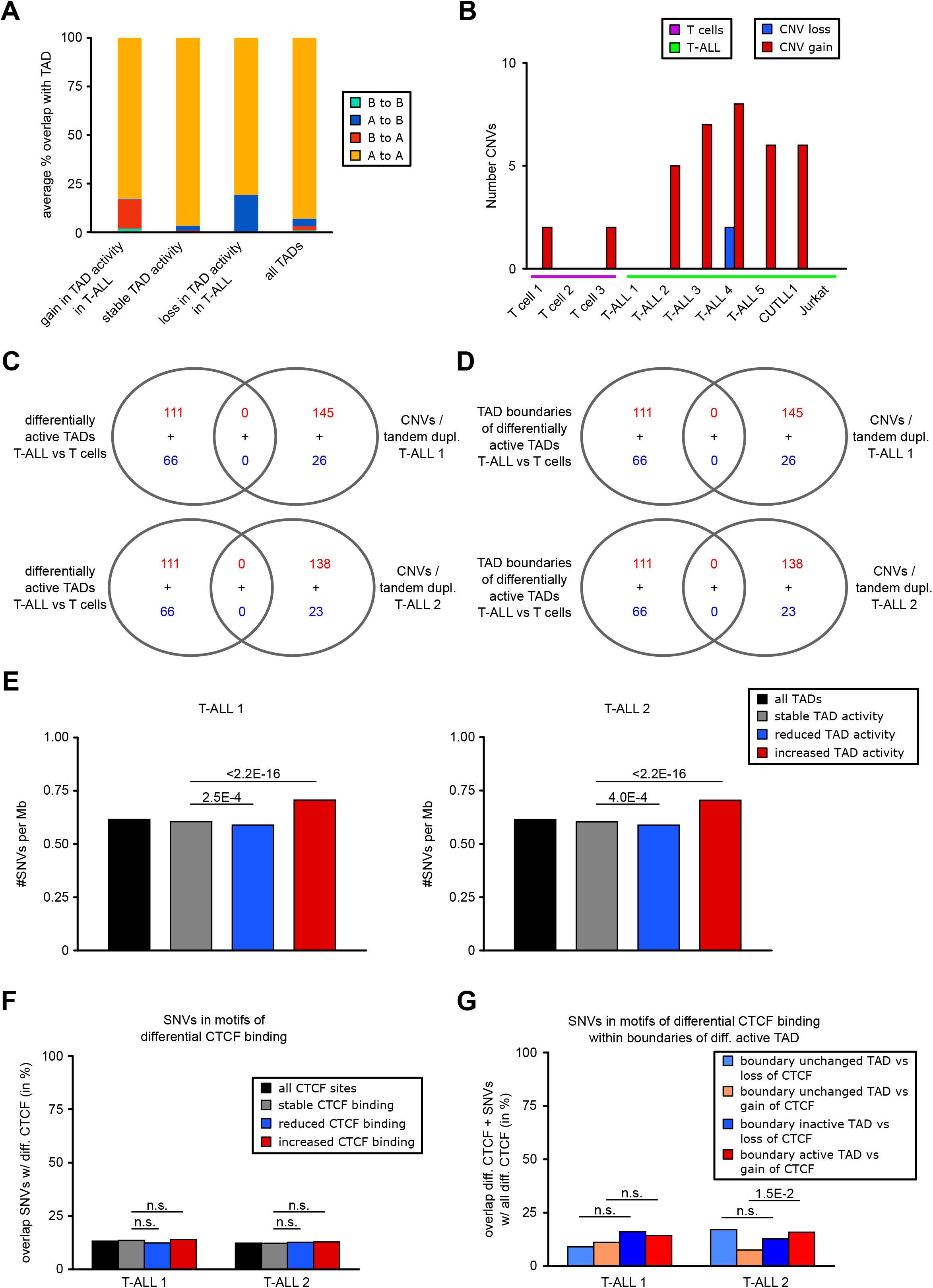
**A)** Copy number variants (CNVs) determined from Hi-C for each sample individually using HiCnv software ^36^. CNV gain was determined if the average estimated copy-number was > 3.5, and CNV loss was determined if the average estimated copy-number was < 1.25. Default setting of chromosome 2 was used as reference chromosome for copy-number estimation. **B)** Average genomic area of TADs (in percent) of differential / stable activity or all TADs overlapping with compartment shifts. Differentially active / stable TADs are defined in Figure 2A; compartment shifts are defined in Figure 1E. **C+D)** WGS detected CNVs (gain / loss) and tandem duplication from overlap with differentially active TADs (C) / boundaries of differentially active TADs (D), both defined in Figure 2A, showed no overlap. Overlap was performed using bedtools intersect, using 1bp overlap between TAD area (C) / TAD boundary extended by 40kb on each side (D) and CNV/tandem duplication. **E)** Integration of SNVs detected from WGS of T-ALL 1 (left) and T-ALL 2 (right) with TAD activity results. SNVs per Megabase (Mb) were counted within genomic areas of all, stably or differentially active TADs. Statistical analysis was performed using two-sided Fisher exact test between numbers of all SNVs overlapping loss/gain of TAD activity and numbers of all SNVs within stably active TADs. **F)** Integration of SNVs with CTCF binding motifs within differential CTCF binding genome-wide. Differential CTCF binding between all profiled T-ALL and T cell samples was determined using DiffBind (FDR < 0.1, log2 fold-change > 1 for increased CTCF binding and log2 fold-change < −1 for decreased CTCF binding in T-ALL; stable CTCF was determined by log2 fold-change > −0.2 and log2 fold-change < 0.2). Statistical analysis was performed using two-sided Fisher exact test between overlap of SNVs with differential CTCF binding and overlap of SNVs with stable CTCF binding. **G)** Integration of SNVs with CTCF binding motifs within differential CTCF binding that overlap with TAD boundaries of differentially / stably active TADs. Differential CTCF binding between all profiled T-ALL and T cell samples was determined using DiffBind (FDR < 0.1, log2 fold-change > 1 for increased CTCF binding and log2 fold-change < −1 for decreased CTCF binding in T-ALL; stable CTCF was determined by log2 fold-change > −0.2 and log2 fold-change < 0.2). Statistical analysis was performed using two-sided Fisher exact test between overlap of SNVs with differential CTCF binding and overlap of SNVs with stable CTCF binding.

**Supplementary Figure 3:**
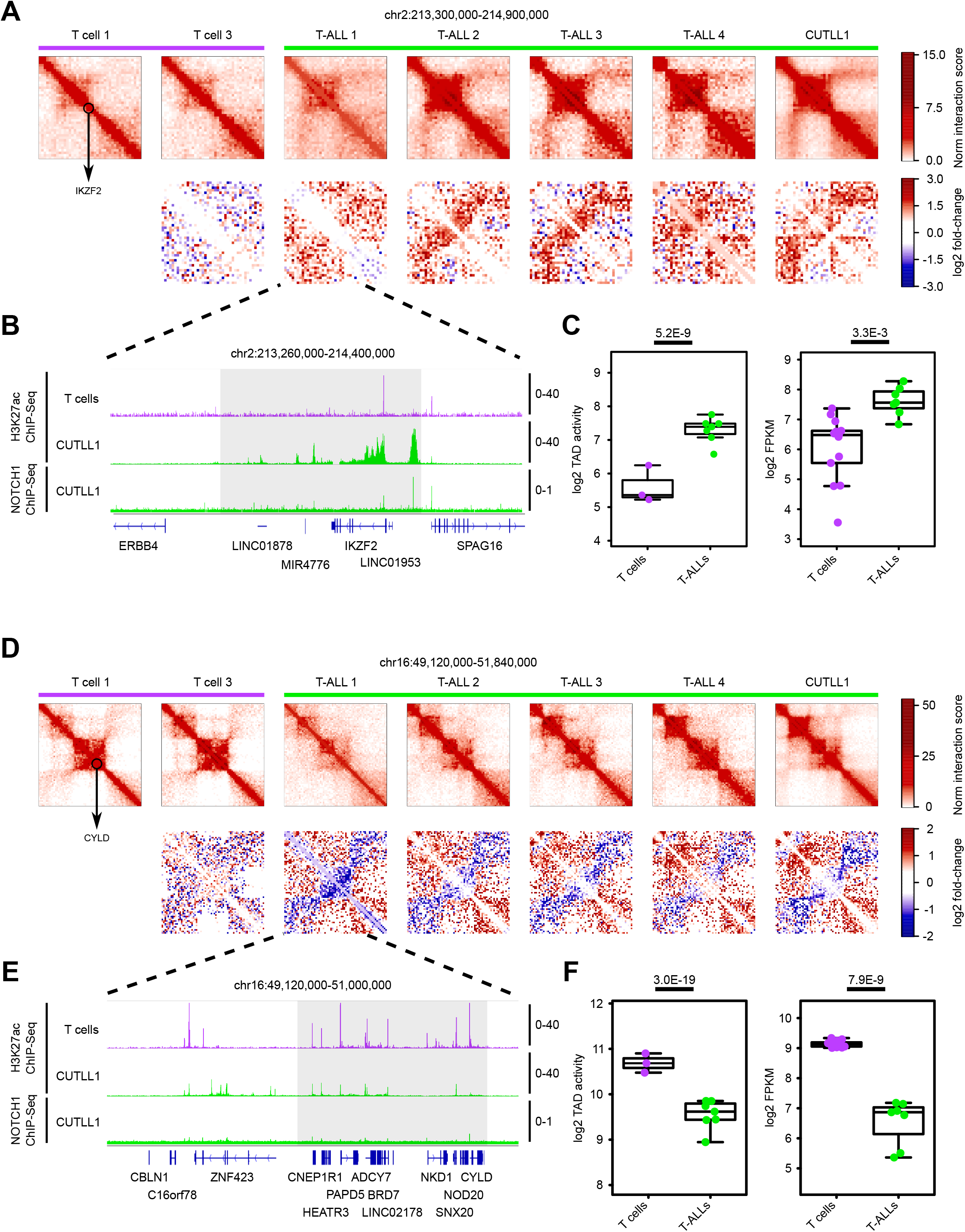
**A)** Hi-C interaction heatmaps (first row) showing the IKZF2 locus (black circle). Second row shows heatmaps of log2 fold-change interactions when compared to T cell 1. **B)** H3K27ac ChIP-Seq tracks for *IKZF2* locus in T cells and CUTLL1, NOTCH1 ChIP-Seq tracks for CUTLL1. Tracks represent fold-enrichment over input where applicable and counts-per-million reads otherwise. **C)** Quantifications for intra-TAD activity (left; as highlighted in A)) and expression of IKZF2 (right). Statistical evaluation for intra-TAD activity was performed using paired two-sided t-test of average per interaction-bin for *IKZF2* TAD between T cells and T-ALL, followed by multiple testing correction (see methods). IKZF2 expression was determined by RNA-Seq and shown as log2 counts per million (CPM) for T cells and T-ALL samples; statistical evaluation was performed using edgeR followed by multiple testing correction. **D)** Hi-C interaction heatmaps (first row) showing the *CYLD* locus (black circle). Second row shows heatmaps of log2 fold-change interactions when compared to T-cell 1. **E)** H3K27ac ChIP-Seq tracks for *CYLD* locus in T cells and CUTLL1, NOTCH1 ChIP-Seq tracks for CUTLL1. Tracks represent fold-enrichment over input where applicable and counts-per-million reads otherwise. **F)** Quantifications for intra-TAD activity (left; as highlighted in D)) and expression of CYLD (right). Statistical evaluation for intra-TAD activity was performed using paired two-sided t-test of average per interaction-bin for CYLD TAD between T cells and T-ALL, followed by multiple testing correction (see methods). CYLD expression was determined by RNA-Seq and shown as log2 counts per million (CPM) for T cells and T-ALL samples; statistical evaluation was performed using edgeR followed by multiple testing correction.

**Supplementary Figure 4:**
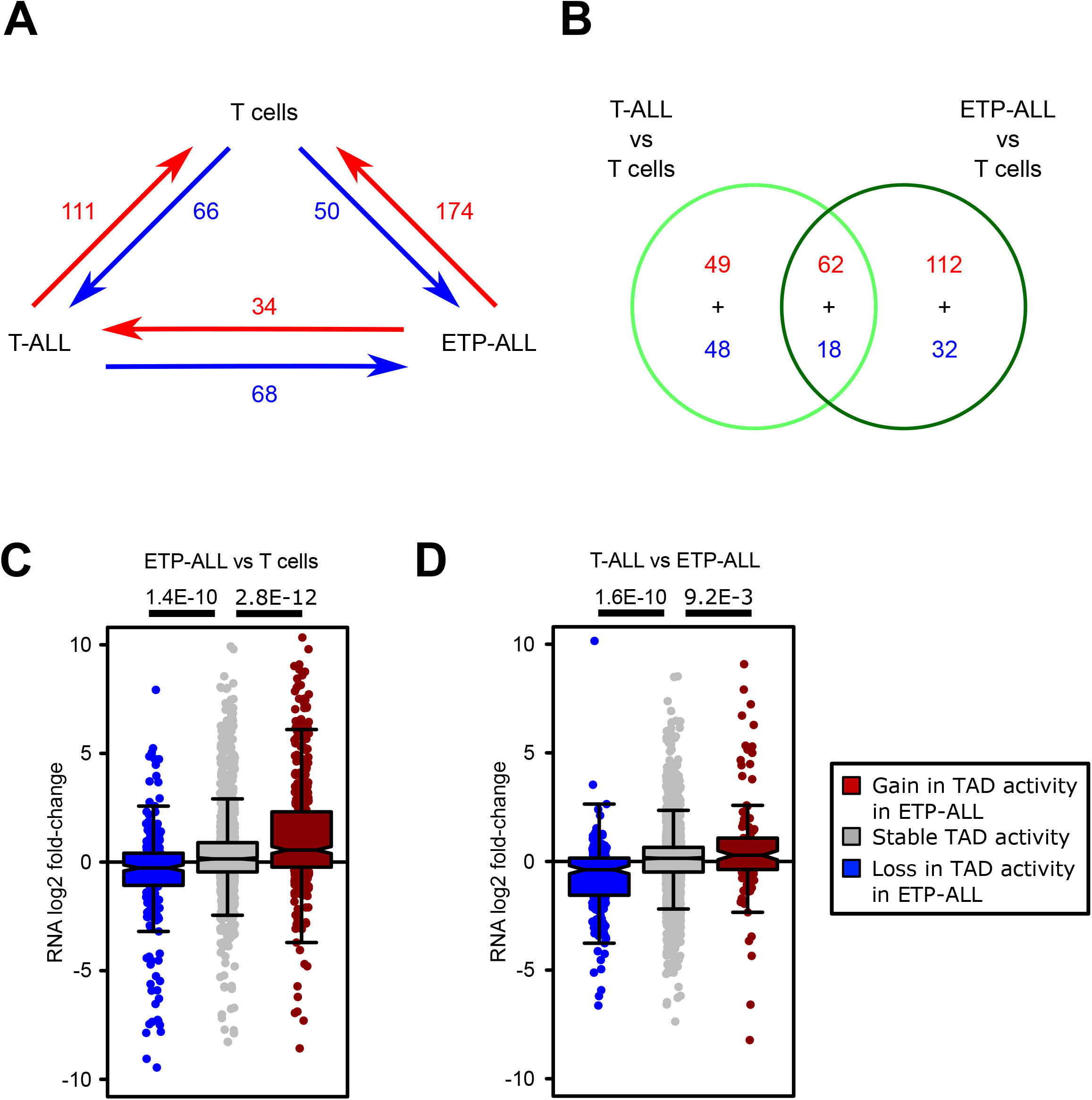
**A)** Comparisons of intra-TAD activity between T cells, T-ALL and ETP-ALL samples. **B)** Overlap of differentially active TADs between the two comparisons of T cells vs T-ALL and T cells vs ETP-ALL, visualized as venn diagram. Red and blue colors correspond to differences as highlighted in A). **C+D)** Integration of RNA-Seq (minimum per-gene expression filter FPKM > 1) within TADs with decreased / increased intra-TAD activity for ETP-ALL vs T cells (C) and ETP-ALL vs T-ALL (D). For each such gene, the respective log2 fold-change in expression between ETP-ALL and T cells (C) / T-ALL and ETP-ALL (D) taken from RNA-Seq is shown. Significant global differences are calculated by an unpaired one-sided t-test comparing genes from TADs with decreased / increased intra-TAD activity with genes from stable TADs, following the hypothesis of a positive correlation between expression and intra-TAD activity changes.

**Supplementary Figure 5:**
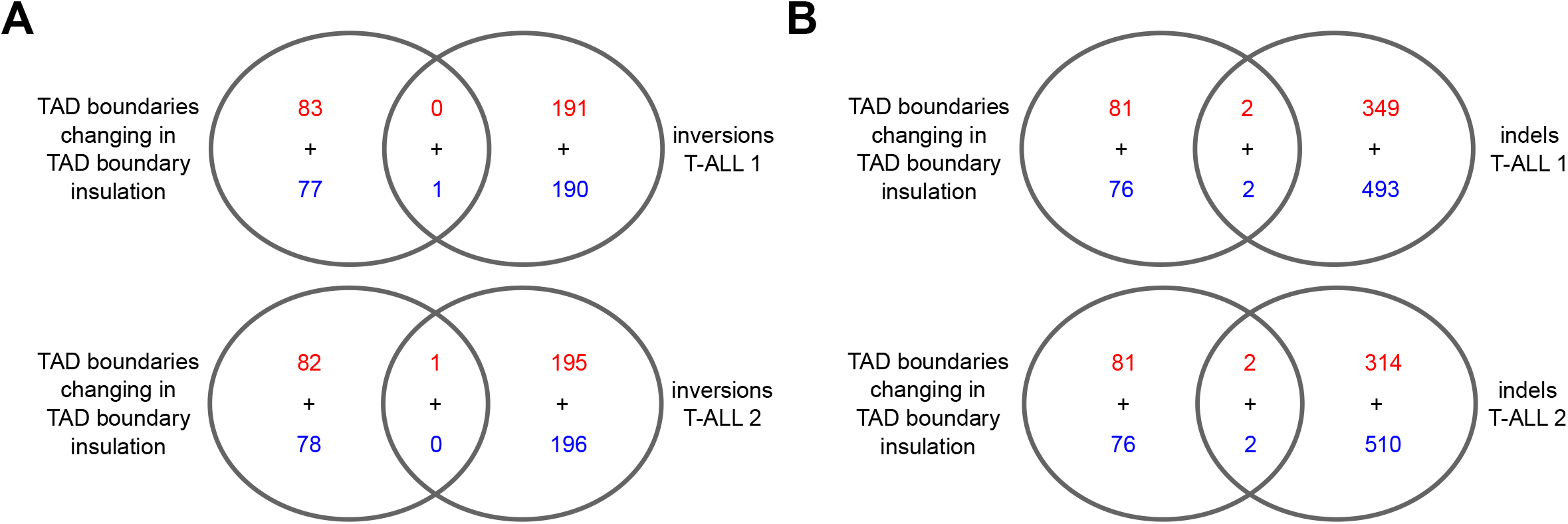
**A+B)** Overlap of TAD boundaries detected as altered in insulation capacity as in Figures 3C and 3D with genomic inversions (A) or insertions/deletions (indels) (B) from WGS of T-ALL 1 (top) and T-ALL 2 (bottom). Overlap was determined by bedtools intersect, using a 1bp overlap.

**Supplementary Figure 6:**
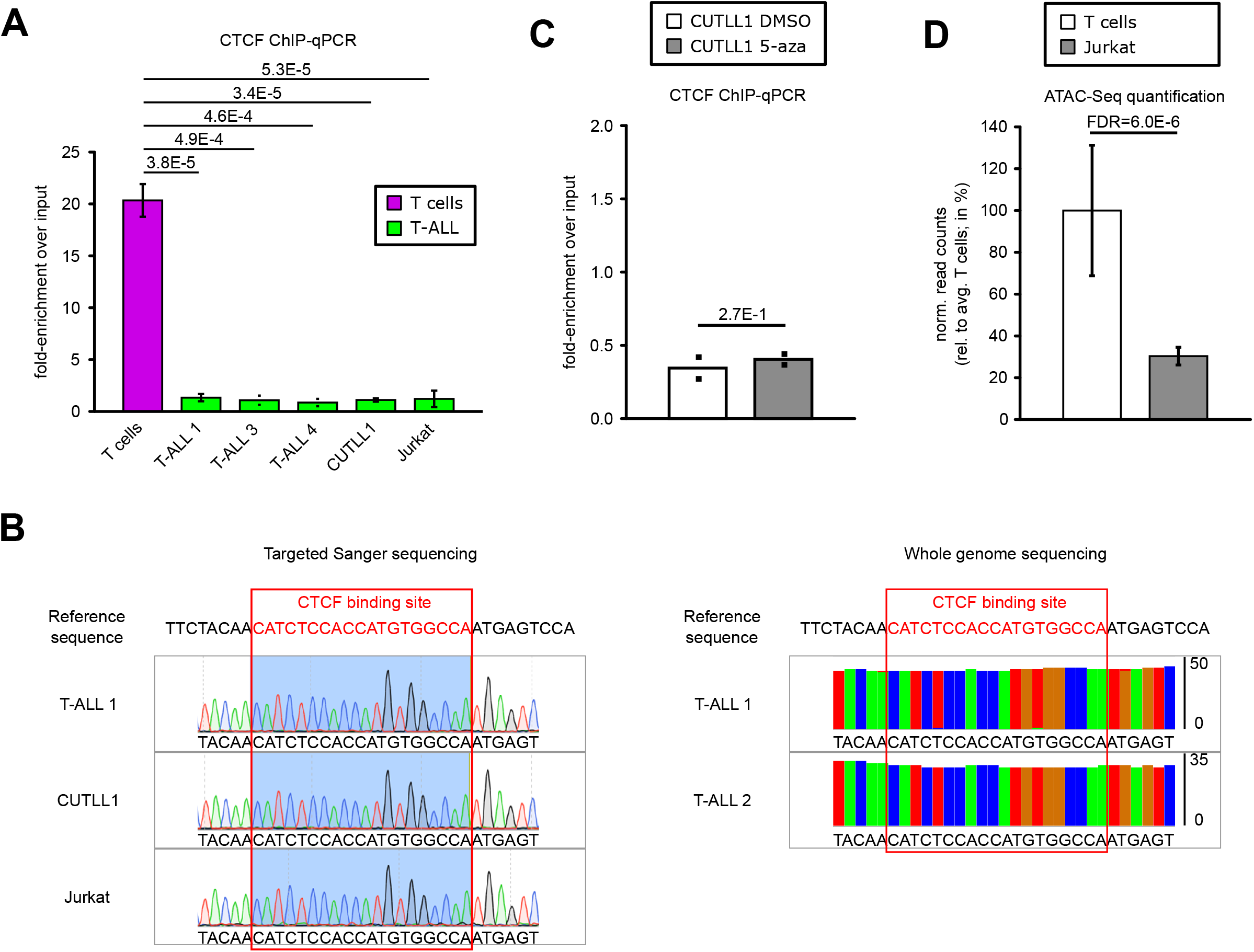
**A)** CTCF ChIP-qPCR of the CTCF binding site in the lost MYC TAD boundary, shown as fold-enrichment over input. Significant differences compared to T cells were calculated with an unpaired one-sided t-test, following the hypothesis of loss of CTCF binding in T-ALL samples as determined from the genome-wide analysis (*n*=3 replicates for T cells, T-ALL 1, T-ALL 2, CUTLL1 and Jurkat; *n*=2 replicates for T-ALL 3 and T-ALL 4). **B)** Targeted sanger sequencing indicates no mutation in T-ALL in the motif of CTCF binding site. Tracks show chromatogram of individual base calls (left). Whole genome sequencing indicates no mutation in T-ALL in the motif of CTCF binding site. Tracks show percent (mis-)matches compared to reference sequence in all reads covering the respective genomic position (right). **C)** CTCF ChIP-qPCR before and after treatment with global DNA-demethylation agent 5-azacytidine. Statistical significance was determined using two-sided t-test (*n*=2 replicates). **D)** ATAC-Seq quantification for T cells and Jurkat for the genomic area covering loss of CTCF binding in the downstream TAD boundary of MYC. Data was normalized to the average T cell signal, shown in percent (*n*=3 replicates). Statistical evaluation was performed using DiffBind, following multiple testing correction.

**Supplementary Figure 7:**
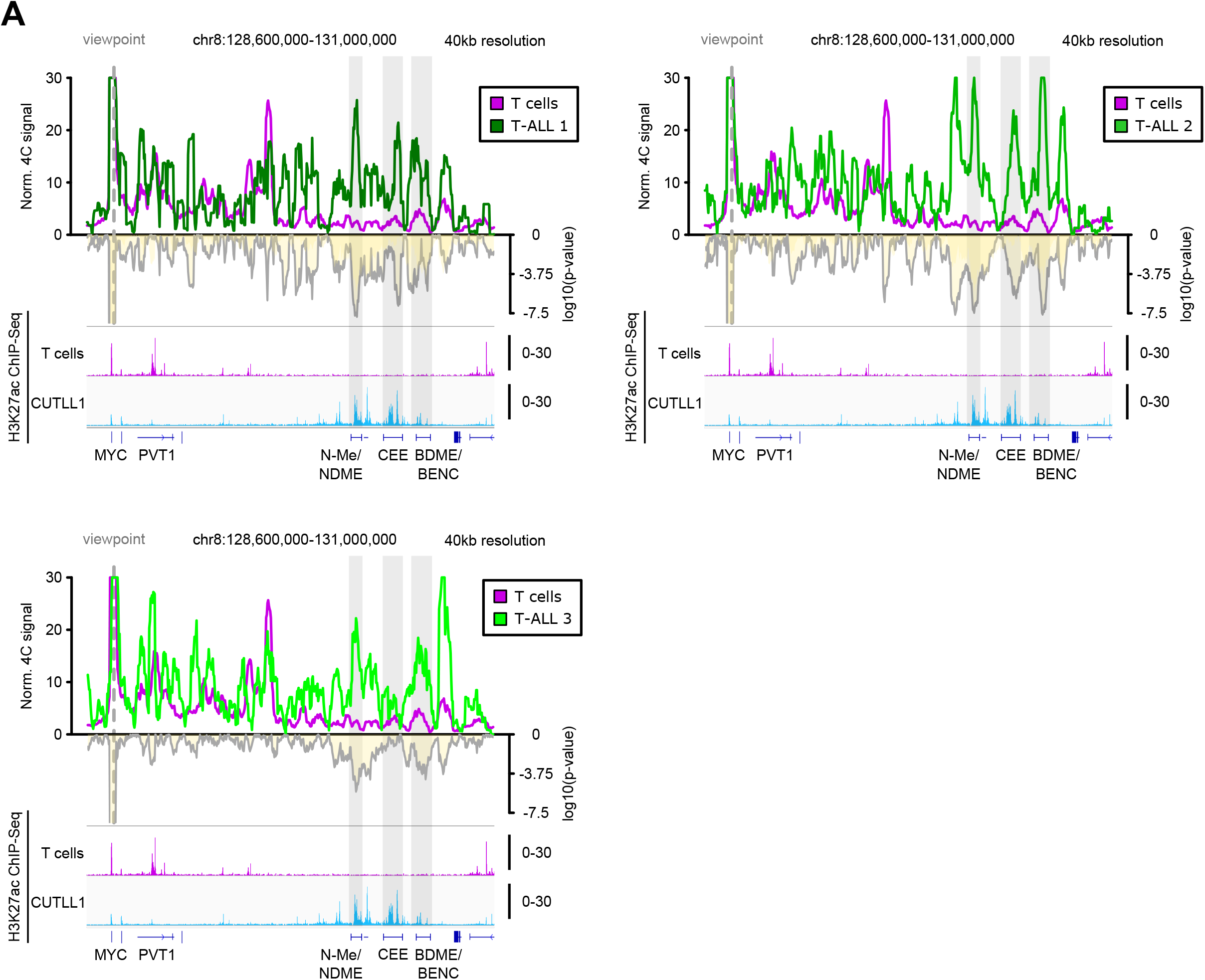
**A)** 4C-Seq analysis using *MYC* promoter as viewpoint. Positive y-axis shows interactions with the *MYC* promoter viewpoint as normalized read counts, while negative y-axis shows significance of differential interactions between T cells and primary T-ALL samples as log(p-value). The three grey boxes highlight three areas of strong H3K27ac signal within the super-enhancer element (N-Me/NDME, CEE, BDME/BENC) that correlate with strong *MYC* promoter interactions. H3K27ac ChIP-Seq tracks for T cells and CUTLL1 are represented below as fold-enrichment over input.

**Supplementary Figure 8:**
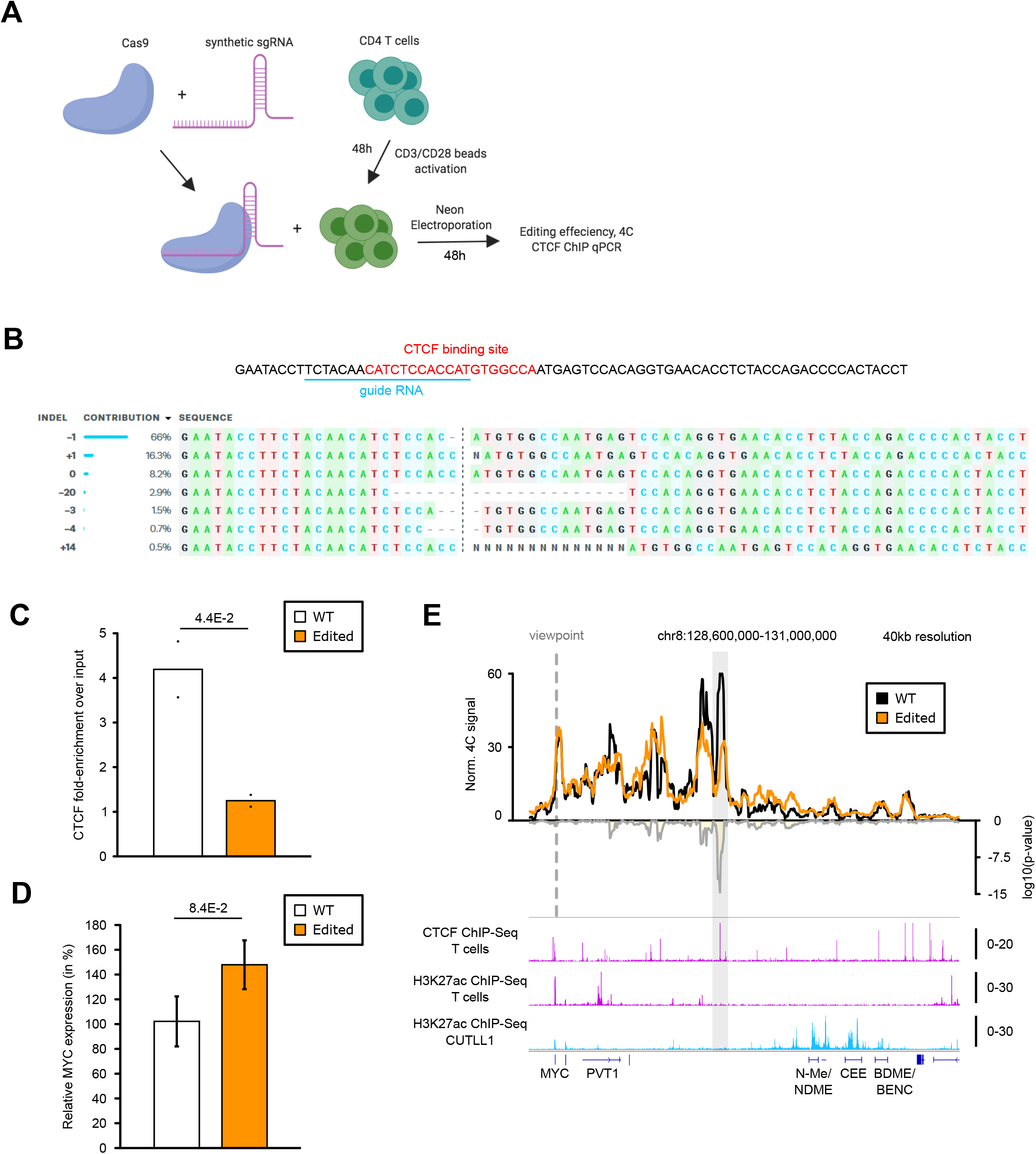
**A)** Schematic of Cas9+Synthetic guide transfection of activated T cells. **B)** Sequence showing CTCF motif in the insulator region in T cells targeted for CRISPR-based deletion. sgRNA targeting sequence within the CTCF motif is highlighted. Sequencing of sgRNA target site indicates various indels along with frequencies observed for each indel. **C)** CTCF ChIP-qPCR validation of reduced CTCF binding in edited T cells compared to unedited T cells (*n*=2 replicates). Statistical significance was determined using unpaired one-sided t-test following the hypothesis that mutation/deletion of the CTCF binding site would lead to abrogation of CTCF binding. **D)** qPCR comparing MYC expression in edited T cells compared to unedited T cells (*n*=3 replicates). Statistical significance was determined using unpaired two-sided t-test. **E)** 4C-Seq analysis using *MYC* promoter as viewpoint in edited and unedited T cells. Positive y-axis shows interactions with the viewpoint as normalized read counts, while negative y-axis shows significance of differential interactions between the two samples as log(p-value) calculated with edgeR function glmQLFTest (*n*=2 replicates). Tracks below show CTCF ChIP-Seq track from CUTLL1 and H3K27ac ChIP-Seq tracks for naïve T cells and CUTLL1 as fold-enrichment over input.

**Supplementary Figure 9:**
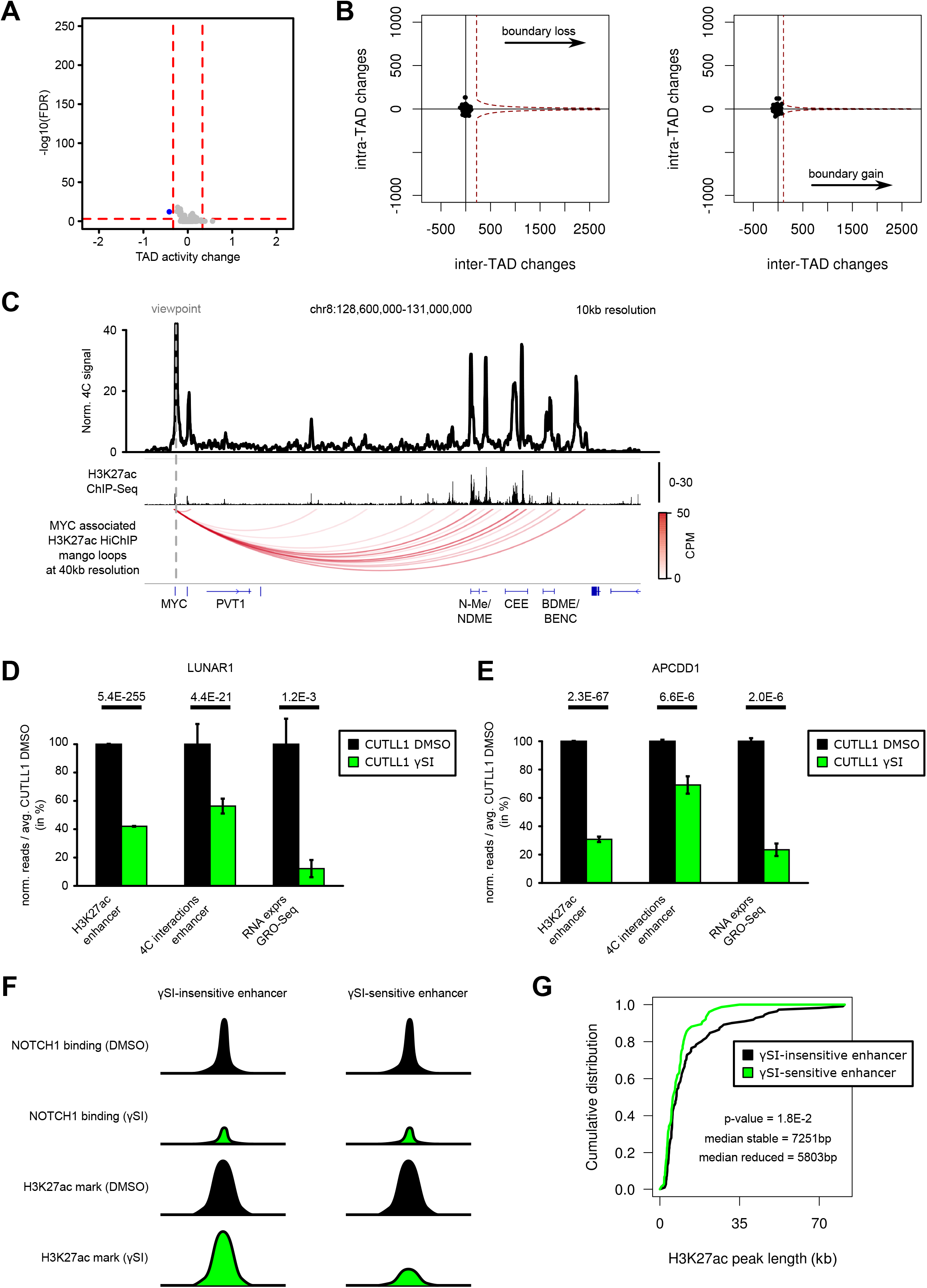
**A)** Volcano plot showing differential intra-TAD activity for the comparison of CUTLL1 treated with either DMSO or γSI. TAD activity changes are highlighted for log2 fold-change of average activity > 0.58 / < −0.58 and with FDR < 0.05. Statistical evaluation was performed using paired two-sided t-test between all per bin-interactions for the same TAD between DMSO and γSI treated cells. **B)** Representation of TAD boundary alteration events (red dots; none identified) among all pairs of adjacent TADs (black dots). Plots depict pair-wise comparisons for TAD boundary losses of adjacent CUTLL1 (untreated, left) TADs and for TAD boundary gains of adjacent CUTLL1 (γSI treated, right) TADs. However, in this analysis, no single TAD boundary alteration was identified reaching the same outlier threshold as the leukemia vs. normal comparison (red dotted lines; taken from Figure 3 C) and D)). **C)** Virtual 4C representation of H3K27ac HiChIP in CUTLL1, using *MYC* promoter as viewpoint (chr8: 128,747,680), showing edgeR-normalized counts-per-million (CPM). H3K27ac ChIP-Seq track for MYC locus in CUTLL1, shown as fold-enrichment over input. Detected significant loops as arc-representation (below) from mango pipeline ^68^ (FDR<0.1; CPM>5). **D)** Quantification of H3K27ac signal (enrichment over input) by ChIP-Seq (left), chromatin interaction of the highest peak by 4C-Seq (center) for the interaction of LUNAR1 promoter with its upstream enhancer element and LUNAR1 expression by GRO-Seq (right). All quantifications are normalized to the respective average T cell signal, shown in percent. Significance of differences was calculated using diffBind (for H3K27ac ChIP-Seq, FDR) and edgeR (for 4C-Seq interactions and GRO-Seq as p-value and FDR respectively). **E)** Quantification of H3K27ac signal by ChIP-Seq (left), chromatin interaction of the highest peak by 4C-Seq (center) for the interaction of *APCDD1* enhancer with the downstream *APCDD1* promoter and APCDD1 expression by GRO-Seq (right). All quantifications are normalized to the respective average T cell signal, shown in percent. Significance of differences was calculated using diffBind (for H3K27ac ChIP-Seq, FDR) and edgeR (for 4C-Seq interactions and GRO-Seq as p-value and FDR respectively). Error bars indicate standard deviation. **F)** Schematic of γSI sensitive and insensitive enhancer. **G)** Comparison of the width of two classes of identified H3K27ac peaks. All peaks are overlapping dynamic NOTCH1 sites, and are of either stable H3K27ac signal (black, *n*=111) or decreased signal (green, *n*=76) as defined in Figure 5A. Significant difference between the distributions is estimated by a two-sided Wilcoxon test.

**Supplementary Figure 10:**
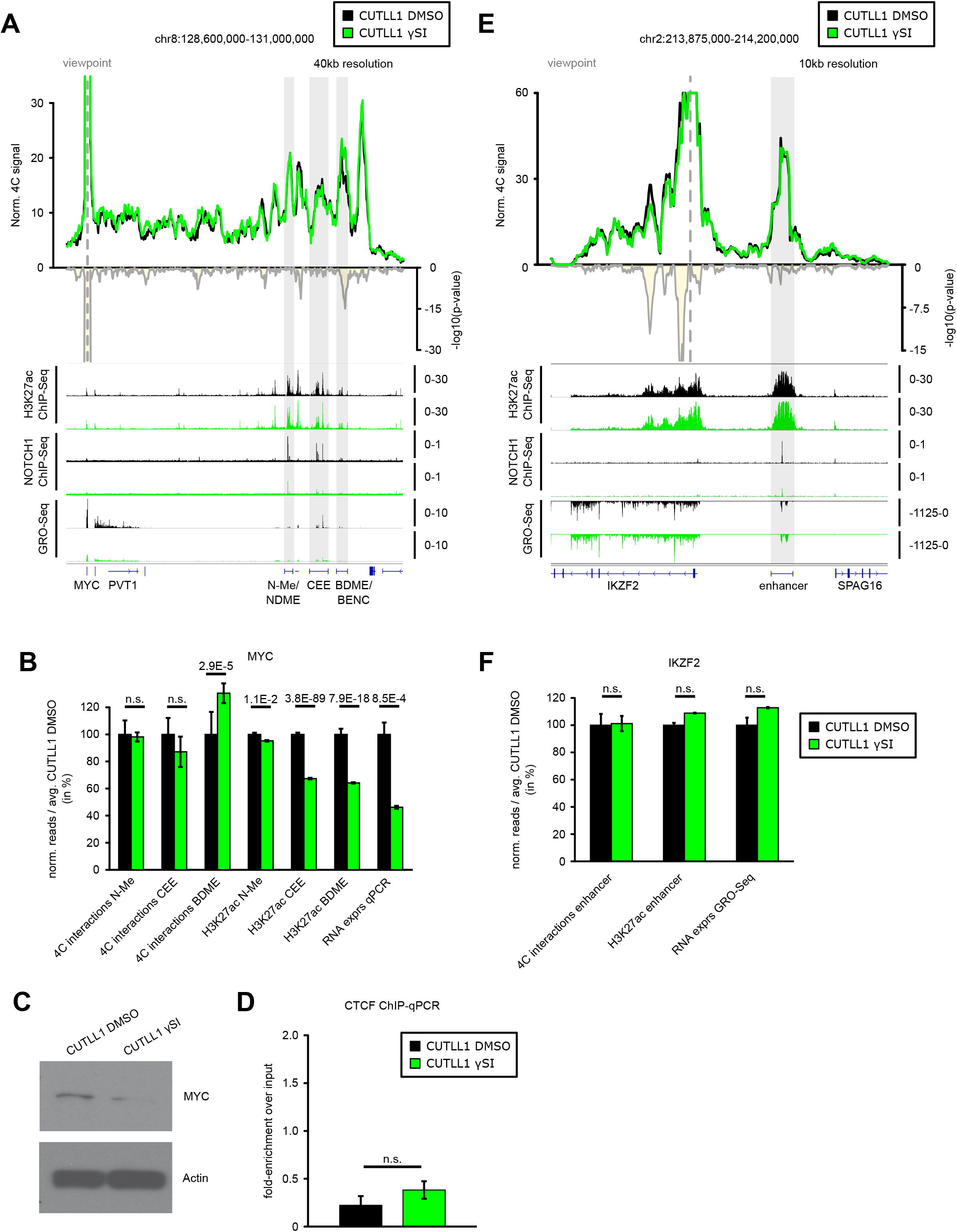
**A)** 4C-Seq analysis using *MYC* promoter as viewpoint after γSI treatment. Positive y-axis shows interactions with the viewpoint as normalized read counts, while negative y-axis shows significance of differential interactions between untreated and γSI treated CUTLL1 as log(p-value) calculated using edgeR function glmQLFTest (*n*=5 for CUTLL1 DMSO; *n*=3 for CUTLL1 γSI). The grey boxes highlight enhancer elements N-Me/NDME, CEE and BDME/BENC. Tracks below show H3K27ac, NOTCH1 ChIP-Seq and GRO-Seq (positive strand only) tracks before and after γSI treatment as fold-enrichment where applicable, and counts-per-million reads otherwise. **B)** Quantification of changes in H3K27ac signal (enrichment over input), chromatin interactions of the highest peaks by 4C-Seq for the interactions of MYC with respective super-enhancer elements and MYC expression after γSI. H3K27ac signal quantification is specific for N-Me/NDME, CEE and BDME/BENC. Interaction changes are measured by centering the 40kb bin on highest peaks within N-Me/NDME, CEE or BDME/BENC elements. MYC expression after γSI treatment was measured by qPCR. All quantifications are normalized to the respective average T cell signal, shown in percent. Significance is shown as false-discovery rate (FDR) for H3K72ac signal change (R package DiffBind), p-value for chromatin interaction change (edgeR function glmQLFTest) or one-tailored t-test for qPCR changes. **C)** Western blot analysis of CUTLL1 cells treated with either DMSO or γSI and immunoblotted with MYC antibody. **D)** 4C-Seq analysis using *IKZF2* promoter as viewpoint after γSI treatment. Positive y-axis shows normalized interaction strength with the viewpoint, while negative y-axis shows significance of differential interactions between untreated and γSI treated CUTLL1 as log(p-value) calculated using edgeR function glmQLFTest (*n*=5 for CUTLL1 DMSO; *n*=3 for CUTLL1 γSI). The grey boxes highlight IKZF2 enhancer element identified by HiChIP analysis. Tracks below show H3K27ac, NOTCH1 ChIP-Seq and GRO-Seq (negative strand only) tracks before and after γSI treatment as fold-enrichment over input where applicable, and counts-per-million reads otherwise. **E)** Quantification of changes in H3K27ac signal (enrichment over input), chromatin interactions and IKZF2 expression after γSI. H3K27ac signal quantification is specific for enhancer highlighted in D). Interaction changes are measured by centering the 40kb bin on the highest enhancer peak. IKZF2 expression after γSI treatment was measured by GRO-Seq. All quantifications are normalized to the respective average T cell signal, shown in percent. Significance is shown as false-discovery rate (FDR) for H3K72ac signal change (R package DiffBind), p-value for chromatin interaction change (edgeR function glmQLFTest) or one-tailored t-test for qPCR changes. **F)** CTCF ChIP-qPCR of lost MYC boundary in DMSO or γSI treated CUTLL1 cells (*n*=3). Significance was calculated using unpaired two-sided t-test.

**Supplementary Figure 11:**
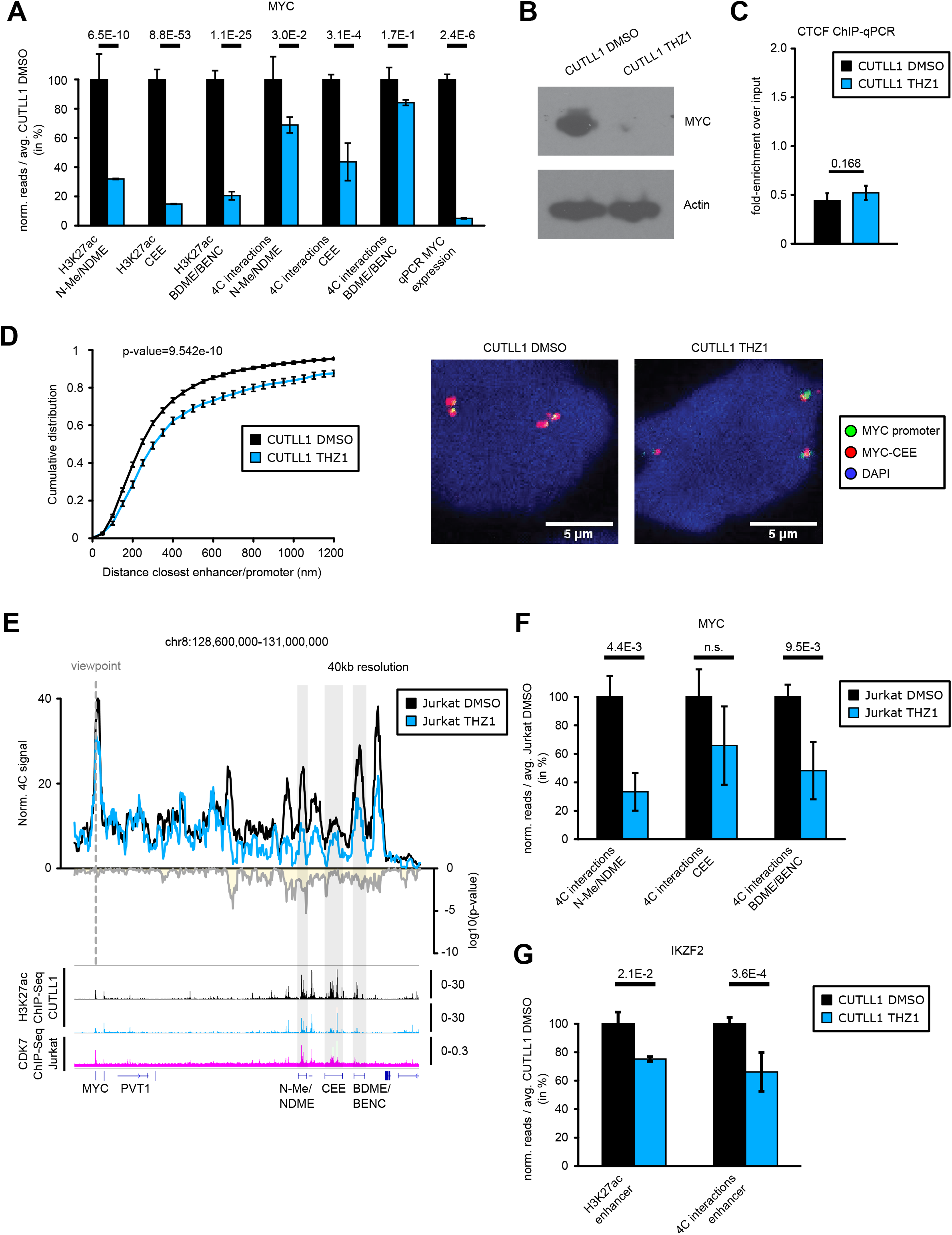
**A)** Quantification of changes in H3K27ac signal (enrichment over input), chromatin interactions and MYC expression after THZ1. H3K27ac signal quantification is specific for N-Me/NDME, CEE and BDME/BENC. Interaction changes are measured by centering the 40kb bin on highest peaks within N-Me/NDME, CEE or BDME/BENC elements. MYC expression after THZ1 treatment was measured by qPCR. All quantifications are normalized to the respective average CUTLL1 DMSO signal, shown in percent. Significance is shown as false-discovery rate (FDR) for H3K72ac signal change (R package DiffBind), p-value for chromatin interaction change (edgeR function glmQLFTest) or two-sided t-test for qPCR changes. **B)** Western blot analysis of CUTLL1 cells treated with either DMSO or THZ1 and immunoblotted with MYC antibody. **C)** CTCF ChIP-qPCR, shown as enrichment over input, of CTCF site in lost boundary in MYC locus in CUTLL1 cells treated with either DMSO or THZ1 (*n*=3). Significance was calculated using unpaired two-sided t-test. **D)** Inter-probe distance between *MYC* promoter and center enhancer element (MYC-CCE) measured by DNA-FISH analysis in CUTLL1 cells treated with either DMSO or THZ1. Statistical difference between distributions of probe distances was calculated using two-sample one-sided Kolmogorov Smirnov test following the hypothesis of increased probe-distance in CUTLL1 cells treated with THZ1 when compared to DMSO. Representative FISH image of CUTLL1 cells treated with either DMSO or THZ1 (right). Probe-pairs CUTLL1 DMSO = 2001. Probe-pairs CUTLL1 THZ1 = 1308. Median distance CUTLL1 DMSO = 264.28µm. Median distance CUTLL1 THZ1 = 321.69µm. **E)** 4C-Seq analysis using *MYC* promoter as viewpoint in Jurkat cells. Positive y-axis shows normalized interaction strength with the viewpoint, while negative y-axis shows significance of differential interactions between untreated and THZ1 treated Jurkat cells as log(p-value) calculated using edgeR function glmQLFTest (*n*=3). **F)** Quantification of changes in chromatin interactions of *MYC* enhancers after THZ1 treatment in Jurkat. Interaction changes are measured by centering the 40kb bin on N-Me/NDME, CEE or the BDME/BENC. Significance is shown as p-value for chromatin interaction changes between DMSO and THZ1 treated cells (edgeR function glmQLFTest). **G)** Quantification of changes in H3K27ac signal (enrichment over input) and chromatin interactions of *IKZF2* enhancer after THZ1 treatment in CUTLL1. All quantifications are normalized to the respective average Jurkat DMSO signal, shown in percent. Significance is shown as p-value for chromatin interaction change (edgeR function glmQLFTest).

## SUPPLEMENTAL TABLE LEGENDS

**Supplementary Table 1:**

References and accession numbers (where applicable) for public Hi-C, ChIP-Seq and RNA-Seq datasets integrated in this study.

**Supplementary Table 2:**

Read alignment statistics for all Hi-C, 4C-Seq, ChIP-Seq, RNA-Seq and GRO-Seq datasets generated within this study.

**Supplementary Table 3:**

List of known driver mutations identified in the primary T-ALL cohort.

**Supplementary Table 4:**

Number of TAD calls per sample using hic-ratio TAD caller ^27^ at 40kb Hi-C matrix resolution and 500kb insulation-window.

## Materials and Methods

### Cell culture

Human cell lines CUTLL1, Jurkat and were cultured in RPMI-1640 media supplemented with 10% fetal bovine serum, penicillin, streptomycin and glutamine. Tissue culture reagents were purchased from Gibco. Naïve CD4 T cells were purchased from Lonza and cultured in X-vivo 15 culture medium (Lonza) substituted with 5% human serum (Gemini Bioproducts) and 10 ng/ml human IL2 at a density of 10^6^ cells per ml.

### Primary T-ALL samples

Primary T-ALL patient samples were collected by Columbia Presbyterian Hospital with informed consent and approved and analyzed under the supervision of the Columbia University Medical Center Institutional Review Board. For expansion of these cells, 1×10^6^ patient cells were transplanted into immunodeficient NOD SCID gamma (NSG) mouse strains via retro-orbital injection as previously performed^1^. Cells collected from the spleen of these primary recipients were used for the *in-situ* Hi-C experiment. All the mouse experiments were performed as per ethical guidelines set by IACUC and NYU.

### *In-situ* Hi-C

Hi-C was performed as described in Rao et al. ^2^ Primary samples have been processed as one replicate and all cell line experiments were processed in two biological replicates each. Briefly, 20 million cells were fixed in 1% formaldehyde for 10 min. Fixed cells were permeabilized in 1ml lysis buffer (10mM Tris–HCl pH 8, 10mM NaCl, 0.2% NP-40, protease inhibitor cocktail (Sigma) for 15 min on ice, spun down (2000 × g, 5 min, 4°C), and the cell pellets were resuspended in 345μl of 1× NEBuffer2 (NEB) per 5 million cell aliquot. 38μl of 1% SDS was added to each aliquot, followed by incubation at 65°C for 10 min. 43μl of 10% Triton X-100 was then added to quench the SDS. To digest chromatin, 400 U of HindIII (NEB) was added per aliquot and incubated at 37°C overnight with continuous agitation (900 rpm). After digestion, restriction sites were filled in with Klenow (NEB) in the presence of biotin-14-dATP (Life Technologies), dCTP, dGTP and dTTP for 2 hours at 37°C. Blunt-end ligation was performed by adding 700µl ligation mix (containing 50U of the T4 DNA ligase (Invitrogen), followed by overnight incubation at 16°C.

The cross-links were reversed by adding 50μl of 10mg/ml proteinase K (Invitrogen) per aliquot and incubated at 65°C for 2 hours, followed by addition of another 50μl 10mg/ml proteinase K and incubated overnight. All the aliquots per replicate were pooled and DNA was extracted by phenol/chloroform extraction protocol. RNA was digested by adding 1μl of 1mg/ml RNase A (Sigma) and incubated at 37°C for 30 min. Biotin was removed from non-ligated restriction fragment ends by incubating 40μg of DNA with T4 DNA polymerase (NEB) for 4 hours at 20°C in the presence of dATP and dGTP. After DNA purification (Amicon Ultra 30K) and sonication (Covaris E220), the sonicated DNA was double-size selected using Ampure XP beads (Beckman Coulter, 0.8 X - 1.1 X). End-repair was performed using T4 DNA polymerase (NEB), T4 DNA polynucleotide kinase (NEB), Klenow (NEB) and dNTPs in 1× T4 DNA ligase reaction buffer (NEB), followed by dATP-addition with Klenow. Biotin-marked ligation products were isolated with MyOne Streptavidin C1 Dynabeads (Life Technologies). Paired-end (PE) adapters (Illumina) were ligated to DNA fragments using 15 U of the T4 DNA ligase (Invitrogen) for 2 hours at room temperature. Bead-bound DNA was amplified with 6 PCR amplification cycles using PE PCR 1.0 and PE PCR 2.0 primers (Illumina). Primary samples T-ALL 2-5, T cell donor 2 and ETP-ALL samples along with CUTLL1 DMSO and THZ1 treated samples were processed with the commercial Arima genomics HiC Kit (https://arimagenomics.com/) and processed according to manufactures guidelines. The concentration and size distribution of Hi-C library DNA after PCR amplification was determined by tapestation (Agilent Technologies), and the Hi-C libraries were sequenced on Illumina Hi-Seq 2500 or Illumina Hi-seq 4000 at 50 cycles.

### ChIP-Seq

ChIP-seq was performed as described previously ^3^. All H3K27ac ChIP-Seq experiments were performed in biological duplicates. CTCF ChIP-Seq experiments for primary samples were performed as biological duplicates. For cell line experiments, we performed five replicates for CUTLL1, three replicates for CUTLL1 γSI experiments and two replicates for CUTLL1 JQ1. For all conditions we created a single input sample. In brief, 5 million cells were fixed in 1% formaldehyde and snap frozen in liquid nitrogen and stored in −80 °C till usage. For Histone chips, 2 million cells were crosslinked as previously described. Nuclei were isolated from the fixed cells using the nuclei isolation buffer (15mM Tris pH 7.5, 60mM KCl, 15mM NaCl, 15mM MgCl2, 1mMCaCl2, 250 mM Sucrose, 1mM DTT and Protease inhibitor). The isolated nuclei were lysed using nuclei lysis buffer (50 mM Tris-HCl (pH 8.0), 10 mM EDTA (pH 8.0) and 1% SDS). This was followed by sonication (30 mins in total) using the bioruptor from Diagenode at high output with 30s ON and 30s OFF cycles. Following sonication to the desired fragment size of 400-600 bp, the sonicated lysate was diluted with nine volumes of IP dilution buffer (0.01% SDS, 1.1% Triton X-100, 1.2 mM EDTA (pH 8.0), 16.7 mM Tris-HCl pH 8.0 and 167 mM NaCl) and magnetic DynaI beads for 1h (preclearing of chromatin). Following preclearing, CTCF was immunoprecipitated with 10 μl of monoclonal rabbit CTCF antibody, clone D31H2 (Cell Signaling 3418) overnight at 4 °C or H3K27ac (Active motif; Catalog no: 39133). The purified ChiP DNA was used to generate sequencing libraries using Hapa Hyper prep kit from Kapa Biosystems. Libraries were sequenced in single-end using Illumina Hiseq 2500 or Illumina Hi-seq 4000 at 50 cycles.

### RNA-Seq

RNA-seq libraries were prepared using NEXTflex Rapid Illumina Directional RNA-seq Library prep kit as per manufacturer’s guidelines. The libraries were sequenced in single-end by either HiSeq 2500 or HiSeq 4000 at 50 cycles.

### 4C-Seq

For LUNAR1 and APCDD1 viewpoints, we have created biological duplicates for all experiments. For MYC viewpoint, we have created five biological replicates for CUTLL1 DMSO, three replicates for CUTLL1 γSI and two replicates for CUTLL1 JQ1 and two replicates for T cells. Edited T cells 2 replicates (5 million each); CUTLL1 DMSO and THZ1 treatment, biological triplicates.

For each replicate, 10 million cells were fixed in 2% formaldehyde and 10% FBS in PBS for 10 min at room temperature. For edited and WT T cells, 5 million cells were used. Crosslinking was quenched with glycine and the 4C-Seq was performed as described previously ^4^. Cells were lysed on ice with 1ml lysis buffer (50mM Tris pH 7.3, 150mM NaCl, 5mM EDTA, 0.5% NP-40, 1% Triton X-100) for 15 min. Nuclei were spun down and resuspended in 360 μl H2O (or frozen). 60 μl of 10X DpnII restriction buffer was added along with 15 μl 10% SDS, and left shaking for 1hr at 37°C followed by addition of 150 μl of 10% Triton X-100 and an additional shaking for 1h at 37°C. 5 μl of undigested control was stored, and nuclei were incubated overnight with 200U of DpnII (NEB, R0543M) restriction enzyme. A fresh 200U of DpnII was added the following morning for 6hrs. Following this, the digestion was checked for completion by running 5 μl of sample in a 1% agarose gel. DpnII was inactivated with 80 μl 10% SDS, and a proximity ligation reaction was performed in a 7ml volume using 4000U T4 DNA Ligase (NEB M0202M). Crosslinks were reversed at 65°C overnight after adding 300 μg Proteinase K. Samples were then treated with 300 μg RNAse A for 45 min at 37°C, and DNA was ethanol precipitated. A 2nd restriction digest was performed overnight in a 500 μl reaction with 50U Csp6l (Fermentas, ER0211). The enzyme was inactivated at 65°C for 25 min, and a proximity ligation reaction was performed in a 14ml volume with 6000 U T4 DNA Ligase. Sample DNA was ethanol precipitated, and purified using the QIAquick PCR purification kit (Qiagen). To generate 4C-Seq library, 1μg of prepared 4C template was amplified 30 PCR cycles per bait per condition (See Supplementary Table for viewpoint sequences) and the amplified fragments were sequenced in Illumina HiSeq 2500 to generate single end reads at 50 cycles.

### HiChIP

HiChIP was performed as previously described^5^ with some modifications. In brief, up to 10 million crosslinked cells were resuspended in 500 μL of ice-cold HiC lysis buffer (10 mM Tris-HCl pH 7.5, 10 mM NaCl, 0.2% NP-40, 1× protease inhibitors) and rotated at 4°C for 30 min. Nuclei were pelleted and washed once with 500 μL of ice-cold HiC lysis buffer. Pellet was then resuspended in 100 μL of 0.5% SDS and incubated at 62°C for 10 min. 285 μL of water and 50 μL of 10% Triton X-100 were added, and samples were rotated at 37°C for 15 min. 50 μL of NEB Buffer 2 and 15 μL of 25 U/μL MboI restriction enzyme (NEB, R0147) were then added, and sample was rotated at 37°C for 2 h. MboI was then heat inactivated at 62°C for 20 min. We added 52 μL of incorporation master mix: 37.5 μL of 0.4 mM biotin–dATP (Jena Biosciences, NU-835-BIO14-S); 1.5 μL of a dCTP, dGTP, and dTTP mix at 10 mM each; and 10 μL of 5 U/μL DNA Polymerase I, Large (Klenow) Fragment (NEB, M0210). The reactions were then rotated at 37°C for 1 h. 948 μL of ligation master mix was then added: 150 μL of 10× NEB T4 DNA ligase buffer with 10 mM ATP (NEB, B0202), 125 μL of 10% Triton X-100, 3 μL of 50 mg/mL BSA (Thermo Fisher, AM2616), 10 μL of 400 U/μL T4 DNA Ligase (NEB, M0202), and 660 μL of water. The reactions were then rotated at room temperature for 4 h. After proximity ligation, the nuclei were pelleted and the supernatant was removed. The nuclear pellet was brought up to 880 μL in Nuclear Lysis Buffer (50 mM Tris-HCl pH 7.5, 10 mM EDTA, 0.5% SDS, 1× Roche protease inhibitors, 11697498001), and sonicated with a Bioruptor 300 (Diagenode) for 8 cycles of 30sec each, on a medium setting. Clarified samples were transferred to Eppendorf tubes and diluted five times with ChIP Dilution Buffer (0.01% SDS, 1.1% Triton X-100, 1.2 mM EDTA, 16.7 mM Tris-HCl pH 7.5, 167 mM NaCl). Cells were precleared with 30 μL of Protein G dynabeads (Life technology #10004D) in rotation at 4°C for 1 h. Supernatants were transferred into fresh tubes and antibody was added (3ug H3K27Ac antibody for 10 million cells) and incubated overnight at 4°C. The next day 30 μL of Protein G dynabeads were added to samples and rotated at 4°C for 2 h. After bead capture, beads were washed three times each with low-salt wash buffer (0.1% SDS, 1% Triton X-100, 2 mM EDTA, 20 mM Tris-HCl pH 7.5, 150 mM NaCl), high-salt wash buffer (0.1% SDS, 1% Triton X-100, 2 mM EDTA, 20 mM Tris-HCl pH 7.5, 500 mM NaCl), and LiCl wash buffer (10 mM Tris-HCl pH 7.5, 250 mM LiCl, 1% NP-40, 1% sodium deoxycholate, 1 mM EDTA). Samples were eluted with 150 μL of DNA elution buffer (50 mM sodium bicarbonate pH 8.0, 1% SDS, freshly made) and incubated at 37°C for 30 min with rotation. Supernatant was transferred to a fresh tube and elution repeated with another 150 μL elution buffer. 5 μL of Proteinase K (20mg/ml) (Thermo Fisher) were added to the 300 μL reaction and samples were incubated overnight at 65°C. Samples were purified with DNA Clean and Concentrator columns (Zymo Research) and eluted in 10 μL of water. Post-ChIP DNA was quantified by Qubit (Thermo Fisher). 5 μL of Streptavidin C-1 beads (Thermo Fisher) were washed with Tween Wash Buffer (5 mM Tris-HCl pH 7.5, 0.5 mM EDTA, 1 M NaCl, 0.05% Tween-20) then resuspended in 10 μL of 2× biotin binding buffer (10 mM Tris-HCl pH 7.5, 1 mM EDTA, 2 M NaCl). Beads were added to the samples and incubated at room temperature for 15 min with shaking. After capture, beads were washed twice by adding 500 μL of Tween Wash Buffer and incubated at 55°C for 2 min with shaking. Samples were then washed in 100 μL of 1× TD Buffer (2× TD Buffer is 20 mM Tris-HCl pH 7.5, 10 mM magnesium chloride, 20% dimethylformamide). After washes, beads were resuspended in 25 μL of 2× TD Buffer, Tn5 (for 50 ng of post-ChIP DNA we used 2.5 μL of Tn5), and water to 50 μL. Tn5 amount was adjusted linearly for different amounts of post-ChIP DNA, with a maximum amount of 4 μL of Tn5. Samples were incubated at 55°C with interval shaking for 10 min. After removing the supernatant 50 mM EDTA was added to samples and incubated with interval shaking at 50°C for 30 min. Beads were then washed two times each in 50 mM EDTA then Tween Wash Buffer at 55°C for 2 min. Lastly, beads were washed in 10 mM Tris before PCR amplification. Beads were resuspended in 25 μL of Phusion HF 2× (New England Biosciences), 1 μL of each Nextera Ad1_noMX and Nextera Ad2.X at 12.5 μM, and 23 μL of water. The following PCR program was performed: 72°C for 5 min, 98°C for 1 min, then cycle at 98°C for 15 s, 63°C for 30 s, and 72°C for 1 min (cycle number was estimated based on the amount of material from the post-ChIP Qubit (approximately 50 ng was run in six cycles, while 25 ng was run in seven, 12.5 ng was run in eight, etc.). Size selection was performed using two-sided size selection with the Ampure XP beads. After PCR, libraries were placed on a magnet and eluted into new tubes. 25 μL of Ampure XP beads were added, and the supernatant was kept to capture fragments less than 700 bp. Supernatant was transferred to a new tube, and 15 μL of fresh beads was added to capture fragments greater than 300 bp. After size selection, libraries were quantified with Qbit and sent for Bioanalyzer to check for the quality and final size of the library. Libraries were sequenced on an Illumina HiSeq4000 platform on PE50 mode.

### *In vitro* drug treatment

CUTLL1 cells were treated with gamma secretase inhibitor (Compound E) purchased from Alexis Bioscience at a 1 μM final concentration. Treatment was performed every 12 hours for 72 hours. THZ1 was purchased from Cayman Chemical (Catalog no: 9002215) and the cells were treated at 100 nM final concentration every 12 h for 24 h. For 5-azacytidine, the cells were treated with 100 nM every day for 3 days (72 h).

### GRO-Seq and library preparation

GRO-seq sequencing were performed in CUTLL1 cells treated with either DMSO or γSI at 1μM for 72h. All experiments were performed in biological duplicates. Gro-seq sample preparation was performed as described previously ^6^. Briefly, nuclei were isolated in swelling buffer (10 mM Tris-HCl pH 7.5, 2 mM MgCl_2_, 3 mM CaCl_2_), lysed twice in lysis buffer (10 mM Tris-HCl pH 7.5, 2 mM MgCl_2_, 3 mM CaCl_2_, 10% glycerol, 0.5% NP-40) and snap-frozen in freezing buffer (50 mM Tris pH 8.0, 40% glycerol, 5 mM MgCl_2_, 0.1 mM EDTA), For run-on reaction, an equal volume of reaction buffer was added to thawed nuclei (10 mM Tris pH 8.0, 5 mM MgCl_2_, 300 mM KCl, 500 μM ATP, 500 μM GTP, 5 μM CTP, 500 μM BrUTP, 1 mM dithiothreitol, 100 U ml^−1^SuperaseIN, 1% Sarkosyl), mixed and incubated at 30 °C for 5 min. The reaction was stopped with Trizol reagent and RNA was phenol/chloroform extracted and ethanol precipitated. RNA was heated in fragmentation buffer (40 mM Tris pH 8.0, 100 mM KCl, 6.25 mM MgCl_2_, 1 mM dithiothreitol), DNAse treated and purified using Zymo RNA Clean & Concentrator (Zymo Research) using the >17nt protocol. Run-on RNA was immunoprecipitated using BSA-blocked BrDU beads (Santa Cruz) in binding buffer (SSPE 0.5X, 1 mM EDTA, 0.05% Tween-20) for 1 h at 4 °C, washed and eluted in elution buffer (5 mM Tris pH 7.5, 300 mM NaCl, 20 mM dithiothreitol, 1 mM EDTA, 1% SDS) at 65 °C for 20 min. Nascent RNA was further phenol/chloroform extracted and sequencing libraries were prepared.

### qPCR

RNA was extracted using the RNeasy Mini Kit using Qiagen kit (Catalog no: 74106) folllowing manufacturer’s guidelines. cDNA was generated using High Capacity RNA-to-cDNA kit from Life Technologies (Catalog no: 4387406) following manufacturer’s guidelines. cDNA was used to perform qPCR using Light cycler 480 SYBR green I Master Mix from Roche Diagnostis (Catalog no: 04887352001). See Supplementary Table for primer sequences. The reactions were run in Roche Light cycler 480 II.

### Sanger sequencing of CTCF site in MYC locus

Genomic DNA from CUTLL1, Jurkat and T-ALL1 were isolated using Qiagen DNeasy kit as per manufacturer’s guidelines. Target locus was PCR amplified using Phusion High Fidelity PCR Master Mix (Thermo Fisher; Catolog no. F531S) using 100 ng genomic DNA as template. Primer sequences are listed in the table below. PCR product was purified using Qiagen PCR purification column and submitted for Sanger sequencing to Genewiz.

### Immunoblotting

CUTLL1 cells treated with DMSO, γSI or THZ1 were pelleted and lysed using RIPA lysis and extraction buffer (Thermo Fisher, Catolog no: 89900). The lysates were boiled with Laemmli buffer, resolved by SDS-PAGE, transferred to PVDF membranes and proteins visualized by immunoblotting. c-MYC (D84C12) rabbit monoclonal antibody was purchased from Cell signaling and anti-actin antibody was purchased from Millipore (Catalog no. MAB1501R)

### CTCF targeting gRNA sequence

The guide RNA target sequence is UCUACAACAUCUCCACCAUG. The guide RNA along with the tracer RNA was purchased as a synthetic guide RNA from Synthego with 2′-O-methyl 3′ phosphorothioate modifications of the first and last three nucleotides.

### Editing of T cells

Naïve T cells were activated with CD3/CD28 beads from Thermo Fisher Scientific (Catalog no: 11161D) for 48 h. Following activation, the CD3/CD28 beads were magnetically removed and 2 million activated T cells were transfected by electrotransfer with either Cas9 (1.5µg) protein and 1µg guide RNA ribonucleoprotein complex or Cas9 (1.5 µg) protein alone for every 200,000 cells using a Neon Transfection system at 1200 V, Width 40 and 1 pulse. Following electroporation, the cells were diluted into culture medium at 10^6^ cells per ml. The electroporation step was repeated after 24 h. 48 h post second transfection, genomic DNA was isolated. Target CTCF region was PCR amplified and subjected to Sanger sequencing. Editing efficiency was computed using the ICE computational program from Synthego.

### High-throughput 3D DNA-FISH

#### Generation of FISH probes

Custom FISH probes targeting MYC promoter and enhancer were designed using the SureDesign custom oligo design tool from Agilent with homology to the regions of interest mined from the hg19 genome build using the default parameters of the SureDesign tool. The MYC promoter probe library targeted a 60 Kb region centered around the promoter whereas the enhancer probe library targeted a 100 Kb region targeting the center enhancer element of the MYC super-enhancer cluster.

#### 3D-FISH experimental protocol

3D-FISH was performed using the Dako FISH Histology accessory kit from Agilent (Catlog no: K579911-5) as per manufacturer guidelines. Briefly, 200,000 cells were cytospun to poly-L-lysine-treated glass slides at 1200 rpm for 5 min. Cells were subsequently fixed for 10 minutes with 4% formaldehyde in PBS at room temperature (RT), followed by membrane permeabilization with 0.5% Triton X-100 in PBS for 20 minutes at RT. The slides were washed once in 1X PBS followed by RNAse treatment (100 µg/ml RNAse A in 2X SSC buffer). The cells were then wash with 2X SSC and dehydrated through alcohol series: 2X 100% ethanol and 2X 70% ethanol, 2 minutes each at RT. The slides were washed with 1X Dako Wash buffer for 5 minutes at RT and treated with 1X Dako pre-treatment solution at 98 °C for 2 minutes and allowed to cool down for 15 minutes at RT. Following pre-treatment, the slides were washed twice with 1X Dako Wash buffer for 3 minutes each at RT. Then the slides were treated with cold pepsin at 37°C for 2 minutes followed by two washes with 1X Dako Wash buffer for 3 minutes each at RT. Then the slides were dehydrated through a series of ethanol washes 70% ethanol, 80% ethanol and 100% ethanol, 2 minutes each at RT. Following the ethanol washes, the slides were air dried and set up for probe hybridization. For each slide 1µl of each probe mixed with 9 µl of IQFISH Fast Hybridization buffer were added, covered with a coverslip and sealed with rubber cement. The slides were incubated at 80°C in a heat block for 10 minutes followed by 90-minute incubation in a hot air oven set at 45°C in dark. Following hybridization, the rubber cement was removed and the slides were washed with 1X Dako stringent wash buffer for 5 minutes at RT immediately followed by a second wash with 1X Dako stringent wash buffer for 10 minutes at 56°C. The stringent washes were followed with two washes of 1X Dako Wash buffer for 3 minutes each at RT. The slides were then dehydrated through a series of ethanol washes 70% ethanol, 80% ethanol and 100% ethanol, 2 minutes each at RT, air dried and mounted with coverslips using immune-mount with DAPI stain.

#### 3D FISH quantification of MYC enhancer-promoter distances

To quantify enhancer-promoter distances from DNA FISH data, we first used AirLocalize (Lionnet et al, Nature Methods 2011) to automatically detect spots in enhancer and promoter channels and estimated their position with subpixel resolution. For enhancers (red channel), the detection parameters were set to σxy = 2.4621, σz = 1.1768 pixels and the intensity threshold to 5000 counts (typical voxel sizes: 73 nm in xy, 360 nm in z). For promoters (green channel), the detection parameters were set to σxy = 2.8801, σz = 1.1776 pixels and the intensity threshold to 4000 counts. Then we filtered the spots by eliminating spots located outside the nucleus or with integrated intensity lower than 1.5e6 counts in red or 0.75e6 counts in green. Next, we computed and plotted the histograms of the distance matrix between spots in the red and green channels. For a perfectly aligned and corrected system, the means of the dx, dy and dz histograms should be 0 in all three directions for there is no reason for enhancers to prefer one relative orientation to promoters than others. We therefore corrected the offsets of the two channels by subtracting the means of the dx, dy and dz histograms from individual coordinate differences. After correction, we computed the distance matrix again. In the distance matrix, we found pairs of spots that are the nearest neighbors to each other, mutual nearest neighbors, which we defined as pairs of enhancers and promoters and built the matrix of their distances. Finally, we plotted cumulative probability distributions of enhancer-promoter distances in the different conditions (N = 30, 23 and 16 image stacks for the CUTTL1-DMSO, CUTTL1-THZ1 and T cells respectively; a typical image contains 10-20 nuclei; probe-pairs T cells = 993, probe-pairs CUTLL1 DMSO = 2001, probe-pairs CUTLL1 THZ1 = 1308). Results were robust to changes in bin size, subsets of images analyzed, or slight changes in imaging conditions, or considering all nearest neighbors rather than only mutual nearest neighbors. Significance for differential co-localization was derived using a Kolmogorov-Smirnow test.

#### T cell donor Information

|  | Age (Years) | Sex |
| --- | --- | --- |
| T_cell_donor | 60 | Male |
| T_cell_donor | 41 | Male |
| T_cell_donor | 63 | Male |
| T_cell_donor | 38 | Male |
| T_cell_donor | 19 | Male |
| T_cell_donor | 60 | Male |

## Bioinformatics analysis

### Hi-C analysis

*In-situ* Hi-C datasets were analyzed with the HiC-bench platform^7^. In short, both datatypes were aligned against the human reference genome (GRCh37/hg19) by bowtie2 (version 2.3.1)^8^ with mostly default parameters (specific settings: --very-sensitive-local --local). For Hi-C, aligned reads were filtered by the GenomicTools^9^ tools-hic filter command (integrated in HiC-bench), which discards multi-mapped reads (“multihit”), read-pairs with only one mappable read (“single sided”), duplicated read-pairs (“ds.duplicate”), read-pairs with a low mapping quality of MAPQ < 20, read-pairs resulting from self-ligated fragments (together called “ds.filtered”) and short-range interactions resulting from read-pairs aligning within 25kb (“ds.too.short”). The reads used for downstream analyses are all accepted intra-chromosomal read-pairs (“ds.accecpted intra”), which were consistently above 25% across all Hi-C samples. The absolute number of accepted intra-chromosomal read-pairs varied between ∼40 and ∼120 million, which in all cases was sufficient to call topologically associated domains (TADs). Interaction matrices for each chromosome separately were created by the HiC-bench platform at a 40kb resolution. Filtered read counts were normalized by a method called “iterative correction and eigenvector decomposition” (ICE) ^10^. To account for variances of read counts of more distant loci, which tend to be less covered the further distant the interacting loci are apart in the genome, we performed distance normalization for each matrix as recently described ^11^.

TADs were called using the algorithm developed within hic-bench ^7^ setting the insulating window to 500kb. The matrix-wide stratum-adjusted correlation score (SCC) was calculated using HiC-Rep ^12^ for each possible pair-wise comparisons of all 14 Hi-C samples. The SCC was first calculated for each pair of chromosome matrices for any possible pair-wise comparison. The final score for a sample-comparison was calculated as the average of all its chromosome scores. Principal Component Analysis (PCA) on Hi-C datasets was performed in R (prcomp, with scale=TRUE and center=TRUE) using the genome-wide Hi-C “ratio” insulation scores for 500kb windows, as defined in Lazaris et al. ^7^. Unsupervised clustering on hic-ratio insulation scores was performed using the R package Mclust version 5.3 (https://cran.r-project.org/web/packages/mclust/index.html), and both EII and VII models found three clusters to be the optimal separation of samples. For visualization of Hi-C data, we created heatmaps for regions of interest using the normalized contact matrices. Heatmaps were generated with the R function image, and color scale was set to the highest normalized score seen in any sample for the particular window. Fold-change heatmaps were generated by calculating the log2 fold-change for each matrix bin vs. T cell 1 sample.

### CTCF & H3K27ac ChIP-Seq analysis

ChIP-Seq datasets were analyzed with the HiC-bench platform ^7^. The ChIP-Seq aligned reads were further filtered by discarding reads with low mapping quality (MAPQ < 20) and duplicated reads using picard-tools (https://github.com/broadinstitute/picard). The remaining reads were analyzed by applying the peak-calling algorithm MACS2 (version 2.0.1) ^13^ with input as control (option -c) wherever applicable. Binding of transcription factor CTCF was determined from narrow-peak calls, while histone-marks were determined from broad-peak calls (option --broad). For differential binding affinity analysis, we ran the R Bioconductor package diffBind with default parameters, which outputs p-value, false-discovery rate and fold-changes of binding affinity for all identified peaks from either sample of any possible pair-wise comparison. For normalization during diffBind, we used the option “method=DBA_EDGER”. For visualization, we generated bigwig tracks (with the help of bedtools version 2.27.1) as fold-enrichment combining all replicates of the actual sample over input wherever applicable using the MACS2 bdgcmp function (with “-m FE”). All bigwig tracks shown were created with IGV (version 2.3.83). CTCF orientation for canonical CTCF binding sites depicted in all tracks with CTCF ChIP-Seq was generated by PWMScan ^14^ (database JASPAR CORE vertebrates; filtered by p-value < 1E-5). Differential binding heatmaps and peak signal quantification were generated with deeptools (version 2.3.3) ^15^ and visualized the 2.5kb up- and downstream of the peak-summit.

### RNA-Seq

RNA-Seq reads were aligned against the human reference genome (GRCh37/hg19) using the STAR aligner (version 2.5.0c)^16^ mostly with default parameters, discarding all non-uniquely aligned reads. Duplicated reads were discarded using picard-tools. For read counting per gene, we used bamutils count of the ngsutils package (version 0.5.7)^17^ on gene annotations from Ensembl V75 in a stranded manner (options -uniq -multiple complete -library RF). Downstream processing was performed in R with the Bioconductor package edgeR (version 3.14.0)^18^ on stranded gene counts, normalizing for intra- and inter-sample variances (edgeR functions calcNormFactors and estimateTagwiseDisp), resulting in counts-per million (CPM) per annotated gene. For cell line data with multiple replicates (CUTLL1 n=3, Jurkat n=2), CPM values were averaged. Differential expression analysis was performed per condition (leukemia vs. normal T cells) with edgeR functions glmQLFit and glmQLFTest.

### GRO-Seq

GRO-Seq reads were aligned against the human reference sequence GRCh37/hg19 using bowtie (version 1.0.0)^8^. All aligned reads were filtered for unique alignment positions (MAPQ > 20). Next, the remaining reads were counted in a stranded manner per annotated gene in Ensembl Genes V75 using bamutils count of the ngsutils package (version 0.5.7; options -uniq -multiple complete -library RF)^17^. We performed normalization using edgeR^18^ (version 3.14.0; functions calcNormFactors and estimateTagwiseDisp), resulting in counts per millions (CPM) per gene after averaging data from replicates. For visualization, we created bigwig tracks per genomic strand using bedtools coverage (2.27.1) after normalizing for sequencing depth and fragment length of 250bp (bedtools coverage option -fs 250). All tracks were visualized with IGV (version 2.3.83).

### 4C-Seq

4C-Seq reads were processed similarly as described in ^19, 20^. First, a reduced genome reference was created for the human reference genome (GRCh37/hg19) by only considering unique sequence fragments from the reference genome sequence that are adjacent to the restriction sites of the restriction enzyme (DpnII) used during the 4C protocol (following the 4C-ker pipeline ^19^). All reads were aligned against this reduced genome reference by bowtie (version 1.0.0) ^8^, only considering uniquely aligned reads. All self-ligated and undigested fragments were removed (following the 4C-ker pipeline). We further validated that all samples had > 0.5 million mapped reads and > 0.5 cis/trans ratio of mapped reads ^20^. Next, we defined successive overlapping windows of different resolutions (10kb and 40kb), and all adjacent windows are overlapping by 90% of their length (9kb and 36kb respectively). We counted uniquely mapped reads for each window per sample and performed normalization with edgeR (leading to CPM per window). This accomplishes a smoothed signal across samples for different sizes of regions to be plotted (approx. 300kb in Figures 5E, 5F, S10E and 6E using 10kb resolution and ∼2MB in Figures 4C, S7, S8E, S10A, 6D and S11E using 40kb resolution). Data from biological replicates were averaged after normalization for visualization. Differential interactions were identified with edgeR (version 3.14.0) functions glmQLFit and glmQLFTest, and log10(p-value) is shown on the negative y-axis of all 4C plots as indicator for the most significant changes. We have not performed multiple testing correction, as each data-point is dependent due to overlapping windows, and would thus potentially lead to a too stringent correction. Quantifications were calculated for the highest single peak (at 10kb resolution for LUNAR1, APCDD1, IKZF2; at 40kb resolution for MYC) within depicted enhancers/promoters in the respective Figures by grey boxes. Normalized 4C signals, as calculated by cpm-function within edgeR, were further normalized against the average control replicates, and shown in percent. Specific p-values shown in Figures were also taken for the same 10kb/40kb bin showing highest 4C signal within respective enhancer/promoter.

### Compartment analysis

Compartment calling was performed using the filtered Hi-C reads of the hic-bench pipeline for all 14 Hi-C samples individually. The “c-score tool” ^21^ was used to determine the A and B compartments at 100kb windows, using information on active chromatin from H3K27ac ChIP-Seq in T cells, CUTLL1 (for T-ALL) and Loucy (for ETP-ALL) to assign A/B to resulting compartment scores. Windows with missing c-score values for at least one sample are removed from the analysis. P-values were calculated using an unpaired two-sided t-test to determine the statistical significance of compartment shifts for the following comparisons: T-ALL vs T cells, ETP-ALL vs T cells and ETP-ALL vs T-ALL samples. After p-value calculation, the mean c-score for all T-ALL, all ETP-TALL and all T cell samples were calculated. Compartment shifts were determined as “A to A” when the mean c-score values for both conditions are > 0.1, “B to B” shift if the mean c-score value for both conditions is < −0.1, and “A to B”/”B to A” shift if the mean c-score value of one condition is < −0.1 and > 0.1 for the other condition (p-value < 0.1).

Unique compartment shifts for either T-ALL or ETP-ALL were identified by combining the results of the above three comparisons. More specifically, an “A to B” shift is considered T-ALL specific if it is identified as an “A to B” shift in the T-ALL vs T cell comparison, but as a “B to A” event in the ETP-ALL vs T-ALL comparison. A “B to A” shift is considered T-ALL specific when it is identified as a “B to A” shift in the T-ALL vs T cell comparison, but as an “A to B” shift for the ETP-ALL vs T-ALL comparison. Similarly, an “A to B” shift is ETP-ALL specific, when it is found as an “A to B” shift for the ETP-ALL vs T cell comparison and an “A to B” shift in the ETP-ALL vs T-ALL comparison; a “B to A” shift is ETP-ALL specific when it is identified as a “B to A” shift in both ETP-ALL vs T cell and ETP-ALL vs T-ALL comparisons.

### Differential TAD activity and data integration

To identify TADs of differential intra-TAD activity, we developed an algorithm to detect statistically significant overall changes between samples of any two conditions (e.g. T-ALL vs. T cells). Firstly, we identified TADs that are common in both conditions. This was done by only considering TADs whose boundaries on either side of the TAD are as close as three bins between the two samples (i.e. 120kb in a 40kb resolution), setting the boundaries of the common TAD to those which yield the largest TAD. We also set a minimum TAD length to 10 bins (400kb). We further removed TADs that fall in the B compartment in both conditions by at least 75% of the genomic TAD area, to avoid minor changes in TAD activity within highly repressed chromatin. This set of common TADs between any two conditions *c*_1_ and *c*_2_ is denoted as *T*. For each interaction bin, we averaged the Hi-C matrix score across conditions (i.e. all T cell, T-ALL or ETP-ALL samples). Next, we performed a paired two-sided t-test on each single interaction bin within each common TAD between the average Hi-C matrix values per condition and calculated the log2 fold-change between the average scores of all interaction intensities within such TADs between the two samples:

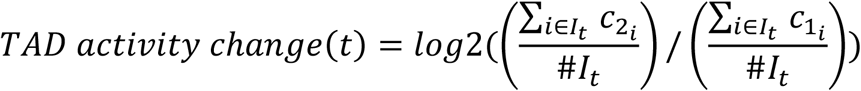

for each *t* ∈ *T*, and *I*_*t*_ being all intra-TAD interactions for TAD *t*.

We also applied multiple testing correction by calculating the false-discovery rate per common TAD (using the R function p.adjust with method=”fdr”). For downstream analyses, we filtered common TADs as differentially active by setting the FDR < 0.1 and absolute log2 fold-change > 0.58. As a negative control group, we defined stable TADs of stable high activity by filtering for TADs with an absolute log2 fold-change < 0.1 and average TAD activity within the top 50% quantile of all TAD activity scores. For downstream CTCF occupancy integration, we extended the TAD boundary for each such identified TAD by 2 bins (80kb) on either side of the boundary accounting for false boundary calls. Changes in CTCF occupancy within these boundaries were defined as the sum of all their log2 FCs taken from the diffBind output, matching the equivalent comparison of T cells vs. T-ALL. Significant changes in global CTCF occupancy within such boundaries were calculated using a one-sided t-test on logFCs from each group (i.e. higher or lower activity in leukemia samples) vs. stable TADs, following the hypothesis of a positive correlation between CTCF binding and TAD boundary strength / TAD activity as recently reported ^22^. Genes (Ensembl V75 annotations; only protein-coding, processed transcripts and lincRNAs with FPKM > 1) were integrated if their promoters were falling within the TADs, extending each TAD by 2 bins (80kb) to either side accounting for inaccurate boundary calls. For each gene, we took the log2 FC from the edgeR output for the respective comparison (T cell vs T-ALL or ETP-ALL vs T-ALL). Significance in global changes of RNA expression was calculated as a one-sided t-test on logFCs from each group (i.e. higher or lower activity in leukemia samples) vs. stable TADs, following the hypothesis of a positive correlation between TAD activity and gene expression.

### Super-enhancer calling and integration

For T cell and CUTLL1 H3K27ac ChIP-Seq data, we applied our standard ChIP-Seq analysis pipeline (https://github.com/NYU-BFX/hic-bench), as described above in detail. Next, we ran ROSE ^23^ with default parameters to define super-enhancers. For each dataset, we have excluded common super-enhancers defined as super-enhancers from both cell-types overlapping by at least 1bp on the genomic coordinates in order to define cell-type specific super-enhancers. We overlapped the remaining cell-type specific super-enhancers with differential active TADs if the overlapping genomic coordinates were larger than 1bp. Enrichment score *ES* of super-enhancers defined as observed over expected overlap was calculated as follows:

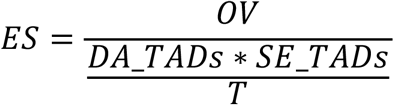

with *SE_TADs* being all TADs containing at least one super-enhancer, *DA_TADs* being all differentially active TADs, *OV* being the intersection of *SE_TADs* and *DA_TADs* and *T* being all TADs the analysis was performed on. Statistically significant enrichment against background (*SE_TADs*) was determined using a two-sided Fisher exact test.

### Compartment shifts within differentially active TADs

We have overlapped compartment information with the differentially active TADs for the T-ALL vs T cell comparison to determine potential compartment shifts within the genomic area of each TAD. Therefore, we have defined “A to B”, “B to A”, “A to A” and “B to B” shifts as described above. To this end, the length of each TAD was determined and the numbers of compartment shifts from any of the previous categories overlapping each TAD were calculated. Then, the percentage of overlap for each TAD was calculated regarding the four compartment shift categories and the average overlap of each category across all TAD categories is shown.

### TAD boundary insulation alterations and differential CTCF integration

We sought to detect more complex changes in chromatin architecture by defining TAD boundary insulation alterations. We separated those into losses and gains of TAD boundaries between normal T cells and leukemia (as depicted in Figure 3A). The analysis was performed in a two-step approach, differentiating between TAD boundary loss and TAD boundary gain as changes resulting from lost versus novel TAD boundaries from the perspective of the leukemia samples, respectively. We thus performed the analysis of identifying insulation changes based on adjacent T cell TADs (yielding TAD boundary losses) and *vice versa* on adjacent leukemia TADs (yielding TAD boundary gains). Thus, for each pair of adjacent TADs for either T cell or T-ALL TAD calls, we determined the interaction strengths of all inter-TAD interactions and intra-TAD interactions for both the two adjacent TADs. The TAD boundary insulation alteration score *BIC* for each pair of adjacent TADs was calculated as *BIC = inter-TAD interactions * max(intra-TAD interactions).* To select the strongest outliers of this analysis as TAD boundary alterations, we took the top 5% quantile of all *BIC* scores between T-ALL and normal T cells. To determine whether these outliers are significant, we performed the same analysis between all three normal T cell donors, applying the same threshold taken from the T cell vs. T-ALL comparison, assuming that there are no severe TAD boundary alterations between any two normal controls. This yielded on average 12 (TAD boundary loss) and 17 (TAD boundary gain) outliers for all three pair-wise comparisons of normal controls, thus we achieved controlling for an average FDR ∼10.77% in T cells vs. T-ALL under the assumption of no boundary insulation changes between normal T cells. For interesting loci, we manually integrated CTCF occupancy and RNA expression changes.

### Genome-wide detection of enhancer activity changes in **γ**SI/THZ1 treated samples

All detected H3K27ac peaks from ChIP-Seq were first overlaid with promoters of annotated genes taken from Ensembl Genes V75. All peaks with a distance of more than 1kb from an annotated promoter (measured from the peak-boundaries) were considered enhancers. Then, we created sets of stable/constant, loss and gain of enhancers in CUTLL1 after γSI or THZ1 treatment using diffBind ^24^ on H3K27ac ChIP-Seq. For stable enhancers, we filtered all peaks with abs(logFC) < 0.2; for reduced/loss of enhancer activity, we filtered all peaks with logFC < - 1.0 / > 1.0 and FDR < 0.05. For γSI-treatment data, we further overlapped all three groups with dynamic NOTCH1-binding sites taken from Wang et al.^25^. Enrichment scores (observed over expected) were calculated similarly as described above, using a two-sided Fisher’s exact test for significance calculation.

### Differential binding analysis using LOLA

In order to define potential co-factors of enhancer/looping activity in γSI-sensitive and insensitive enhancers (Suppl. Figure 9F), we used LOLA ^26^. To this end, we downloaded the LOLA database (http://databio.org/regiondb) and kept ChIP-Seq data from T-ALL related cell lines (Jurkat, CUTLL1 or HPB-ALL), that displayed at least 3000 peaks. We are representing the results as percent overlap between ChIP-Seq peaks and γSI-sensitive / insensitive genomic locations. Statistics for differences between γSI-sensitive and insensitive enhancers was calculated using a two-sided Fisher exact test.

### HiChIP data analysis and loop calling

H3K27ac HiChIP data in CUTLL1 was processed with the hic-bench platform similarly as described above for Hi-C data. We have used output of filtered/accepted intra-chromosomal reads, and performed mango ^27^ to identify significant loops at a 40kb resolution. The trajectories of each matrix bin of the HiChIP data onto both anchors on the diagonal were overlaid with peaks identified from H3K27ac ChIP-Seq in CUTLL1, requiring a minimal overlap of 1bp between a HiChIP-bin and a ChIP-peak. Only loops that were supported by a ChIP-peak in at least one anchor were kept for further analyses. We then applied sequencing-depth normalization (CPM) per replicate followed by a statistical approach described in mango, which employs a binomial test in each diagonal of the counts-matrix up to a maximum distance of 2Mb. High-confidence HiChIP loops were identified by FDR < 0.1 and requiring a minimum CPM > 5 per loop. We have only kept loops that contain an H3K27ac peak outside any annotated promoter in one anchor and an annotated promoter in the other anchor, thus defining promoter-enhancer loops for downstream Hi-C integration analyses.

### Hi-C analysis for **γ**SI/THZ1 treated cells using HiChIP defined enhancer-promoter interactions

Next, to investigate the involvement of changes in enhancer H3K27ac signal in nearby gene expression and loop formation upon γSI/THZ1 treatment in CUTLL1, we integrated Hi-C data with promoter-enhancer loops. To this end, we first identified candidate interactions of promoter-enhancer pairs using loop calling from H3K27ac HiChIP data in CUTLL1, as described above. We further took these specific promoter-enhancer pairs and calculated changes in Hi-C connectivity, using the normalized contact matrices at 40kb resolution. We calculated log2 fold-changes between DMSO and γSI/THZ1 treatment matrices after averaging Hi-C matrices across replicates in each condition. Global loss/gain of interactions upon γSI/THZ1 treatment was depicted by a one-sided t-test comparing all groups vs. the stable H3K27ac group, following the hypothesis of a positive correlation between promoter-enhancer looping and enhancer activity.

### Integration of GRO-Seq data with findings from combined H3K27ac ChIP and HiChIP analysis

For all genes connected with nearby differential/stable enhancers (categorized by ChIP-Seq as described above) detected from HiChIP, we investigated expression of such genes before treatment, after treatment and after 1, 2, 3, 4, 5, 6, and 10 hours post drug “wash off”. We are representing the median FPKM across all genes (FPKM > 1) of a respective enhancer-promoter loop category per time-point. The genome-wide trend of reduced expression for specific H3K27ac categories was determined by a one-sided t-test comparing expression with all genes within stable H3K27ac enhancer-loops, following the hypothesis of a positive correlation between expression changes and looping/enhancer activity.

### WGS analysis and integration with TADs and CTCF binding

Whole-genome sequencing and subsequent data analysis in primary T-ALL samples was performed by GeneWiz (https://www.genewiz.com/). In short, copy number variants were called using Canvas version 1.3.1 and SVs were called using Manta version 0.28.0. Results of CNVs/tandem duplications, other SVs or SNVs were overlapped with genomic areas of (differentially active) TADs or TAD boundaries expanded by 1 bin (40kb) in each direction using bedtools intersect and a minimum overlap of 1bp. Overlap of SVs or SNVs with CTCF binding information was performed by first overlapping differential CTCF peak calls with CTCF motif information derived from PWMScan (database JASPAR CORE vertebrates; filtered by p-value < 1E-5), and then with SVs or SNVs using bedtools intersect and a minimum overlap of 1bp. Significance of overlaps was calculated using two-sided Fisher exact test.

### CNV calling from genome-wide Hi-C data

We have used HiCnv ^28^ in order to detect copy-number variations from all 14 Hi-C samples conducted within this study (excluding cell line data with drug treatments). We have used default parameters, with frag_limit=150 (in the script run_HiCnv.pl) as suggested by authors for Arima Hi-C (combination of frequent 4bp-cutting enzymes) and frag_limit=500 (in the script run_HiCnv.pl) as suggested by authors for Hi-C using HindIII (infrequent 6bp-cutting enzyme). Resulting copy-number variant bins were merged if adjacent genomic bins had the same predicted copy number variant score, and CNVs were called if any such merged bin had a detected CNV of > 3.5 (CNV gain) or < 1.25 (CNV loss).

#### Primer sequences

##### qPCR primers

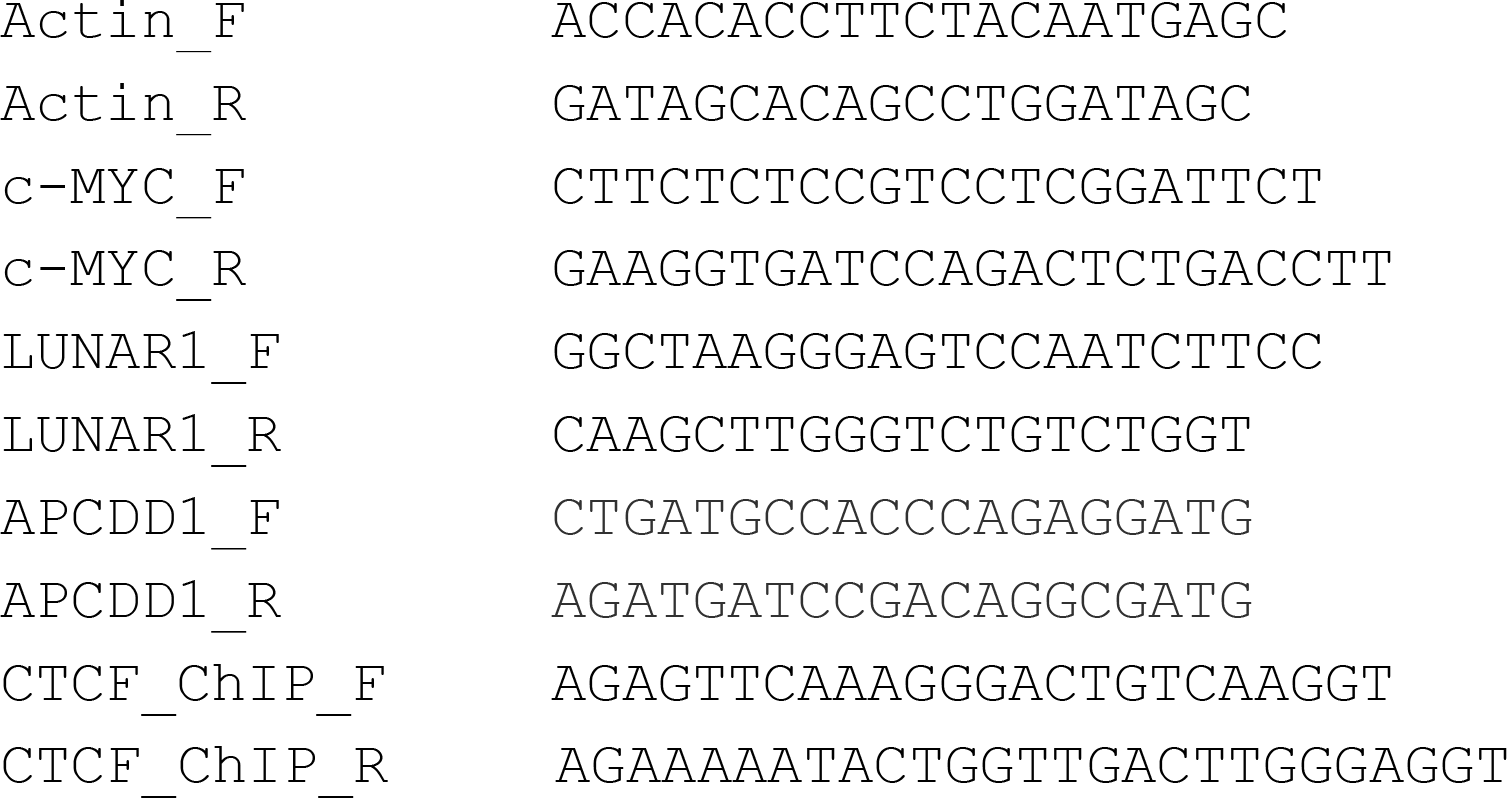

##### 4C primers

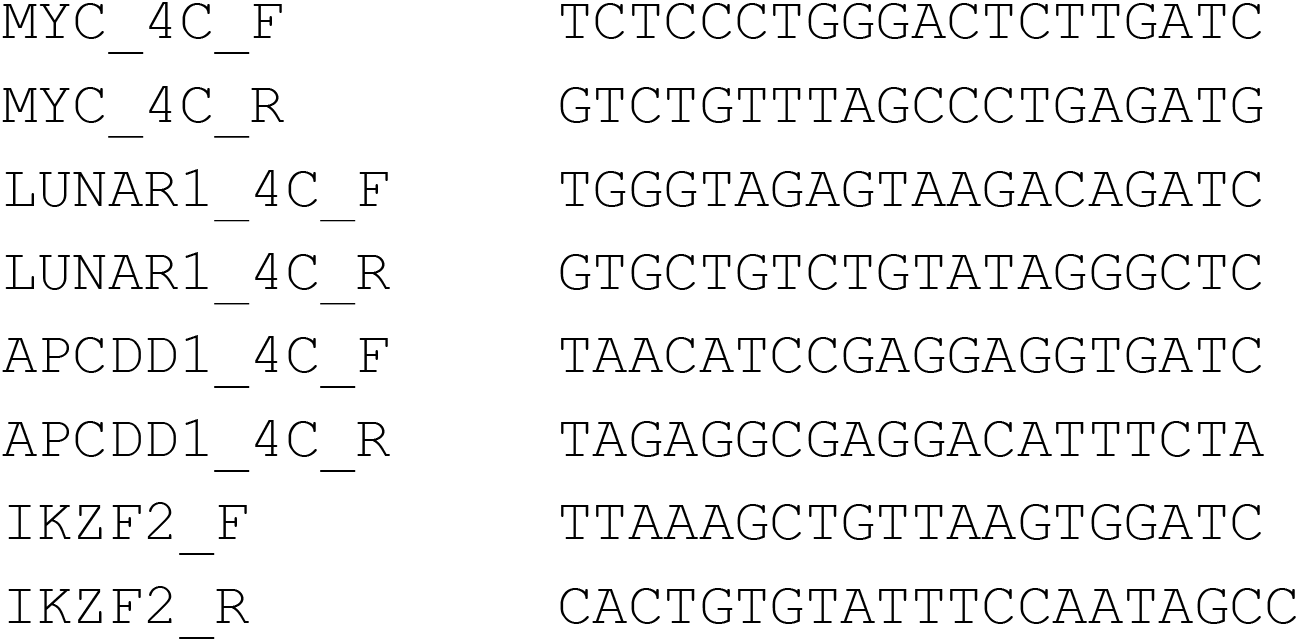

##### Genomic PCR

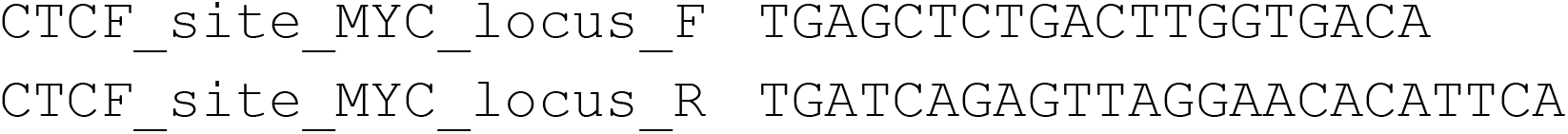

